# Inferring phenotypes of single cells based on the expression profiles of phenotype-associated marker genes in bulks and single cells

**DOI:** 10.1101/2024.12.20.629842

**Authors:** Yin He, Rongzhuo Long, Li Zhao, Xiaosheng Wang

## Abstract

Single-cell transcriptomes is advantageous in characterizing cell types and states, but not sufficient in describing cell phenotypes due to small cohorts in single-cell datasets. Thus, inferring single-cell phenotypes with the leverage of bulk data is practical. Here we proposed an algorithm for Single Cells’ Phenotype Prediction based on the expression profiles of phenotype-associated marker genes in bulks and single cells (ScPP). ScPP first analyzes bulk data to identify phenotype-associated marker genes. Next, ScPP evaluates the enrichment scores of the phenotype-associated marker gene sets in single cells using the AUCell algorithm. The single cells with the phenotype are defined as the intersection of the single cells with top α ranks according to the phenotype-associated AUC values and the single cells with bottom α ranks according to the opposite phenotype-associated AUC values. Finally, all single cells are determined as phenotype+, phenotype- or background. We demonstrated that ScPP could effectively recognize cell subpopulations with specific phenotypes, including tumor malignancy, estrogen receptor status, microsatellite instability, copy number variation, survival and immunotherapy response. Compared to the established algorithms (Scissor and scAB), ScPP displays more excellent predictive performance and needs less running time. Thus, ScPP is an effective and competitive tool for inferring cell phenotypes.

## Introduction

Single-cell transcriptomes reflect the gene expression profiles of single cells, which have been widely used to characterize cell types, states and subpopulations in heterogeneous tissue ecosystems^1^. Specifically, compared to bulk transcriptomes, single-cell transcriptomes gain a huge advantage in investigating the extensive heterogeneity of malignant tumors as well as the tumor microenvironment^2^. Currently, many algorithms have been developed for cell type identification, such as CHETAH^3^, SingleR^4^, scmap-cell^5^ and SciBet^6^, to name a few, and the most commonly used approach for defining cell types based on single-cell transcriptomes is measuring the relative expression abundance of cell-type-specific marker genes^7,8^. Nevertheless, there are limited methods for identifying cell subsets that drive specific phenotypic or clinical features, such as clinical outcomes and therapy response in cancer, although this kind of methods are valuable in identifying cell subpopulations associated with disease diagnosis, prognosis and treatment. Furthermore, most single-cell datasets involve small cohorts that hampers the investigation of the association between cell subpopulations and phenotypic features. By contrast, bulk transcriptomes often involve large cohorts, such as the data from The Cancer Genome Atlas (TCGA)^9^ and International Cancer Genome Consortium (ICGC)^10^, both of which harbor more than 10,000 cancer specimens. Therefore, combining bulk and single-cell transcriptome data may avail complementary advantages in exploring disease phenotypes and heterogeneity.

There are two main research directions in integration of bulk and single-cell data. One is to infer bulk information with the leverage of single-cell data, such as inferring the cellular composition of tumors with reference to single-cell transcriptomes^11,12^. Another is to infer single cell information with the leverage of bulk data, such as inferring phenotypic features of single cells with reference to bulk transcriptome and phenotype data^13,14^. Overall, there are abundant investigations in the former direction^11,15–20^, while the investigations in the latter direction remain insufficient. Scissor^13^ and scAB^14^ are two algorithms recently developed to predict phenotypic features of single cells from single-cell RNA sequencing (scRNA-seq) data by using phenotypic information from bulk data. Scissor was developed using a sparse regression model^13^, and scAB used a knowledge- and graph-guided matrix factorization model to infer single cells’ phenotypes^14^. Apparently, these algorithms further identification of phenotypes beyond cell types in single-cell data analysis. However, owing to the complexity of these algorithms, it is difficult or inefficient to implement these tools in many cases with limited computing resources.

Inspired by the prior methods for cell phenotype and type identification, here we proposed an algorithm for Single cells’ Phenotype Prediction (ScPP) based on the expression profiles of phenotype-associated marker genes in bulks and single cells. By analyzing simulated and real data, we demonstrated that ScPP was capable of effectively recognizing cell subpopulations with phenotypic features of interest, such as malignancy, estrogen receptor (ER) status, microsatellite instability (MSI), copy number variation (CNV), survival prognosis and immunotherapy response. Compared to Scissor and scAB, ScPP displayed more superior performance in identifying phenotype-related cell subpopulations. Furthermore, ScPP gains a significant advantage over Scissor and scAB for its easy and fast implementation in large-scale scRNA-seq datasets. Thus, ScPP is an effective and competitive tool for inferring cell phenotypes and types.

## Methods

### ScPP algorithm

To infer phenotypes of single cells from scRNA-seq data, ScPP requires three types of data as input, including single-cell transcriptomes, bulk transcriptomes, and phenotypic features in bulk data. The phenotypic features can be categorical variables, continuous variables, or clinical survival. For binary variables, such as therapy response versus nonresponse, ScPP first analyzes bulk data to identify significantly upregulated genes in one class versus another class using a threshold of Student’s *t* test adjusted *P* value (false discovery rate (FDR)) < 0.05 and mean expression fold change (FC) > 1.5. The genes significantly upregulated in one class are defined as its marker genes or signatures. Next, ScPP evaluates the enrichment scores of marker gene sets in single cells from single-cell transcriptome data using the AUCell algorithm^21^. Further, ScPP ranks single cells based on their AUC values output by AUCell. Suppose we denote the set of single cells as S1 situating at the upper α of the rank based on the class A’s marker gene set-induced AUC values, and the set of single cells as S2 situating at the bottom α of the rank based on another class’ marker gene set-induced AUC values; ScPP defines the intersection of S1 and S2 as class A+ single cells. Finally, all single cells are determined as class A+, class A-, or background. Class A+ indicates the single cells having the phenotype of class A, class A-the single cells lacking the phenotype of class A (or having an opposite phenotype), and background the single cells predicted as neither Class A+ nor Class A-. For continuous variables and survival variables, ScPP follows the same procedure as above to identify phenotype-associated single cells after attaining related maker genes. For continuous variables, ScPP uses the Pearson or Spearman correlation to identify their marker genes (FDR < 0.05; |correlation coefficient| > 0.2); for survival variables, ScPP utilizes the univariate Cox regression analysis to identify the marker genes significantly correlated with patients’ survival (FDR < 0.05). Here α is a tunable parameter ranging from 20% to 45% (default value: 20%). All results inferred by ScPP that are shown in this paper were obtained using the default α value, unless otherwise specified. A schematic illustration of the ScPP algorithm is shown in Fig. 1. ScPP has been developed into an R package publicly available in GitHub (https://github.com/WangX-Lab/ScPP).

**Fig. 1.**
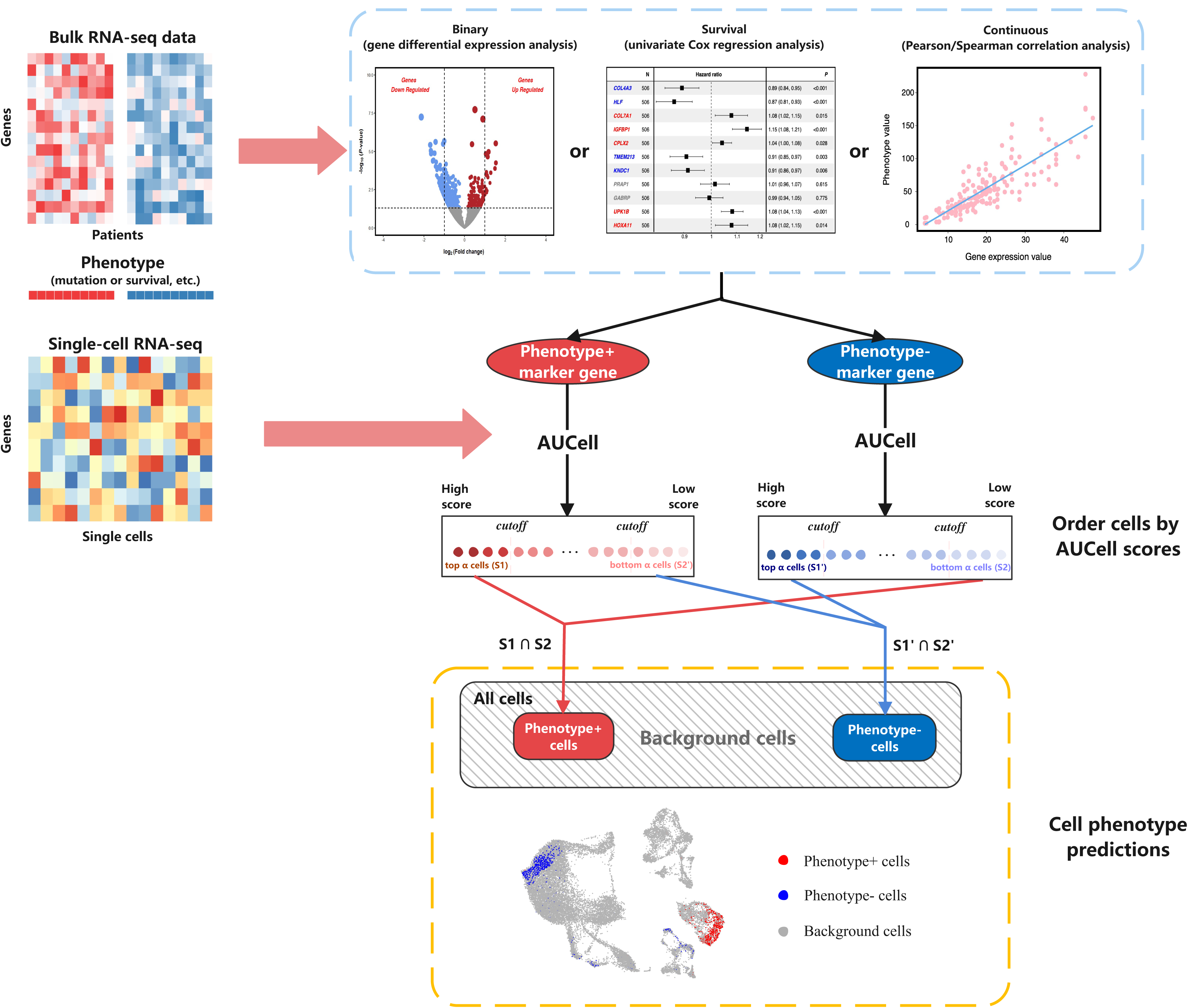
Overview of the ScPP algorithm.

### Datasets

We downloaded bulk transcriptomes (RNA-seq, RSEM normalized), somatic mutations and CNVs, and clinical data for three TCGA cancer types (lung adenocarcinoma (LUAD), breast invasive carcinoma (BRCA) and skin cutaneous melanoma (SKCM)) from the Genomic Data Commons (GDC) data portal (https://portal.gdc.cancer.gov/). The TCGA data for colon and rectal cancer (CRC) were downloaded from the University of California Santa Cruz Xenabrowser (UCSC-Xena)^22^ (https://xenabrowser.net/datapages/). Moreover, we downloaded two BRCA bulk data, namely METABRIC^23^ from cBioPortal (https://www.cbioportal.org/) and GSE113184^24^ from the NCBI Gene Expression Omnibus (GEO) (https://www.ncbi.nlm.nih.gov/geo/). We created a melanoma bulk transcriptome dataset by combining six bulk RNA-seq datasets (PRJEB23709^25^, phs000452^26^, Nathanson^27^, GSE115821^28^, GSE91061^29^ and GSE78220^30^) and correcting for batch effects using the “ComBat()” function in the “sva” R package^31^. This dataset, termed melanoma-Combulk, was the gene expression profiles in 427 melanoma patients treated with immune checkpoint inhibitors (ICIs), including 158 responders and 269 non-responders. Here we analyzed six scRNA-seq datasets, including an LUAD dataset from a previous publication^32^ and five datasets from GEO (GSE149614^33^ for hepatocellular carcinoma (HCC), GSE123813^34^ for squamous cell carcinoma (SCC), GSE176078^35^ for BRCA, GSE115978^36^ for melanoma, and GSE132465^37^ for CRC). The cell annotation and phenotype information in GSE123813 was obtained from an associated publication^38^. A summary of these datasets is shown in Supplementary Tables S1&S2.

### Data simulation

To test the ScPP algorithm, we generated three simulated datasets, termed SD1, SD2 and SD3. SD1 was generated from GSE149614 for identifying malignant and non-malignant cells in HCC; SD2 was created from GSE123813 for predicting responsive and non-responsive T cells in SCC in the immunotherapy setting; and SD3 was produced from a published LUAD scRNA-seq dataset^32^ and TCGA-LUAD for inferring malignant and non-malignant cells in LUAD.

To be specific, to generate SD1 and SD2, we first randomly selected 1,000 cells from the real scRNA-seq datasets, with 500 cells being phenotype-positive (phenotype+) and the other 500 cells phenotype-negative (phenotype-). Next, with the createFolds function in the caret R package, the scRNA-seq dataset composed of the 1,000 selected cells was randomly divided into halves according to the phenotypic information, of which one half was treated as the simulated single-cell dataset for phenotype prediction (termed sscPr), and another half was used for constructing a pseudobulk dataset (termed sscPb). To construct the gene expression profiles in a pseudobulk sample with a specific phenotype, we randomly selected 500 cells with replacement having the phenotype in sscPb and took the average expression of a gene in the 500 cells as its expression in the pseudobulk sample. This procedure was repeated for 100 times to generate a pseudobulk transcriptome dataset containing 50 phenotype+ and 50 phenotype-samples. For SD3, we randomly selected tumor and normal samples with replacement from TCGA-LUAD samples to construct the simulated bulk transcriptome data, which had identical numbers of tumor and normal samples as TCGA-LUAD; meanwhile, we randomly selected 500 malignant and 500 non-malignant cells from the LUAD scRNA-seq dataset^32^ according to the annotation information to construct the simulated single-cell dataset for phenotype prediction. In the data simulations, we set different seed numbers for each simulation to generate 100 different combinations of simulated single-cell and bulk datasets. A summary of the simulated datasets is shown in Supplementary Table S1.

### Prediction performance evaluation

We reported precision or positive predictive value (PPV), negative predictive value (NPV) and balanced accuracy to evaluate the performance of algorithms, which are defined as follows:

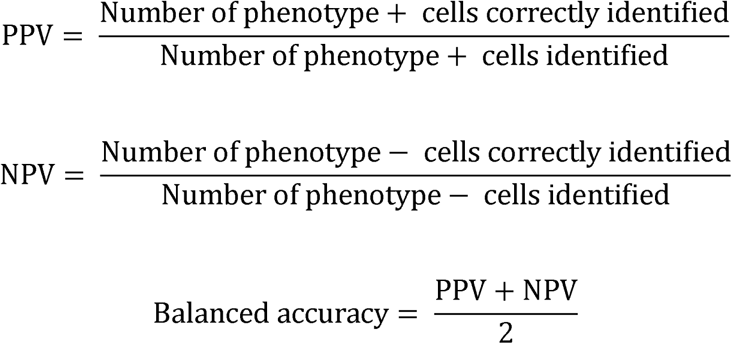

### Data preprocessing

For bulk RNA-seq data, all gene expression values were log_2_(x+1) transformed before subsequent analyses. For scRNA-seq data, we used the Seurat^39^ R package (version 4.3.0) to conduct data preprocessing. First, we used the NormalizeData function to normalize single-cell expression values and FindVariableFeatures to detect highly variable genes across single cells. Based on the highly variable genes, we scaled the data by ScaleData and performed principal component analysis by RunPCA. After that, we utilized FindNeighbors to calculate the similarities between cells based on the first twenty principal components and FindClusters to identify cell clusters. Finally, we performed dimensional reduction and cell visualization using RunUMAP.

### Identifying differentially expressed genes (DEGs)

In bulk datasets, the DEGs were identified using a threshold of Student’s *t* test adjusted *P* value (FDR) < 0.05 and FC > 1.5. The FDR was calculated by the Benjamini-Hochberg method^40^ for adjusting for *P* values in multiple tests. In scRNA-seq datasets, the DEGs were identified by using the FindMarkers function in the Seurat R package with the default of Mann-Whitney *U* test.

### Gene set enrichment analysis

We used the AUCell^21^ R package (version 1.20.2) to evaluate the enrichment scores of gene sets in single cells. First, we employed the AUCell_buildRankings function to compute gene expression rankings of each cell with the expression matrix and default parameters. Based on gene expression rankings, we obtained the area-under-the-curve (AUC) values of each cell by the AUCell_calcAUC function. The AUC values represent the enrichment levels of gene sets in single cells. For bulk data, we evaluated the enrichment scores of gene sets in bulk samples using the single-sample gene-set enrichment analysis (ssGSEA)^41^. The ssGSEA algorithm is a non-parametric method to define a gene set enrichment score per sample as the normalized difference in empirical cumulative distribution functions of gene expression ranks inside and outside the gene set. We performed ssGSEA using the GSVA R package (version 1.46.0) by setting the method = “ssgsea”. In addition, we utilized the website tool GSEA (https://www.gsea-msigdb.org/gsea/msigdb/index.jsp) and gseGO function in the clusterProfiler^42^ R package (version 4.6.2) to perform gene set function and enrichment analysis.

### Cell-cell communication network

We utilized the CellChat^43^ R package (version 1.6.1) to demonstrate the potential regulatory relationships between cells. In detail, we used the function createCellChatto create the object and preprocessed the object by the subsetData function with default parameters. Then, the communication probability was inferred by the computeCommunProb and computeCommunProbPathway functions. Finally, the netVisual_circle, netVisual_bubble and netAnalysis_signalingRole_heatmap functions were used for network visualization.

### Survival analysis

We conducted survival analyses using the survfit function in the survival R package (version 3.5.5). Kaplan-Meier curves were used to display the differences of survival time between different groups of cancer patients, and the log-rank test was utilized to assess the significance of survival time differences. Furthermore, we constructed univariate Cox proportional hazard models using the coxph function in the survival R package to assess the association between clinical features and patients’ survival prognosis. This analysis identified the significant prognostic factors as the input of the multivariable Cox proportional hazard model. The two-tailed Wald test was utilized to assess the association between variates and survival time.

### Identifying T cell subtypes

We used Seurat to cluster T cells with the FindClusters function under the resolution = 0.3. Based on the marker gene expression profiles of T cell subtypes^44^, we identified T cell subtypes.

### Evaluating tumor mutation burden (TMB), intratumor heterogeneity (ITH) and CNVs

TMB was defined as the total number of somatic mutations (excluding silent mutations) in the tumor. We used the DEPTH^45^ algorithm to evaluate ITH. The inferCNV^46^ R package was utilized to estimate initial CNVs for each region of chromosomes. For each cell, its inferCNV-inferred CNV values were re-standardized to the range [-1, 1]. The CNV score of each cell was the quadratic sum of CNV ^47^.

### Comparisons between ScPP and other algorithms

We compared the performance of predicting cell subpopulations related to specific phenotypes between ScPP and two other algorithms, including Scissor^13^ and scAB^14^. The performance included power, accuracy, stability and efficiency. The accuracy included PPV, NPV and balanced accuracy. The stability was measured by the median absolute deviation (MAD) of the accuracies under different parameter settings. The efficiency was evaluated by algorithm’s running time. We ran both algorithms with their default parameters.

### Statistical analysis

In comparisons of two classes of normally distributed data, we used Student’s *t* tests; otherwise we used Mann-Whitney *U* tests. We employed the Spearman or Pearson method to evaluate the correlation between two variables and reported the correlation coefficient and *P* value. To correct for *P* values in multiple tests, we used the Benjamini-Hochberg^40^ method to calculate the FDR. We utilized hypergeometric test (hyper function with parameter “lower.tail = F”) to estimate whether the results inferred by ScPP are due to chance. We performed all statistical analyses in the R programming environment (version 4.2.1).

## Results

We evaluated the performance of ScPP in three simulated datasets (SD1, SD2 and SD3) and six real datasets. To demonstrate the superiority of ScPP over other algorithms for inffering single cell phenotypes, we compared ScPP with two recently published algorithms: Scissor^13^ and scAB^14^, in prediction accuracy, stability and efficiency.

### ScPP shows high prediction accuracy and stability in simulated datasets

We first evaluated the performances of ScPP in a series of simulated datasets using the default parameter (α = 0.2). In the three simulated datasets (SD1, SD2 and SD3), ScPP consistently achieved high PPV, NPV, balanced accuracy and stability. Among 100 simulations, the minimum PPV was 0.938, 0.946 and 0.968, respectively; the minimum NPV was 0.913, 1 and 0.989, respectively; and the minimum balanced accuracy was 0.950, 0.973 and 0.984 in SD1, SD2 and SD3, respectively (Fig. 2a). Moreover, ScPP showed high prediction stability, as evidenced by that: (1) the MAD of PPV among 100 simulations was 0.019, 0 and 0.01, respectively; (2) the MAD of NPV among 100 simulations was 0.021, 0 and 0, respectively; and (3) the MAD of balanced accuracy among 100 simulations was 0.011, 0 and 0.005 in SD1, SD2 and SD3, respectively (Fig. 2b).

**Fig. 2.**
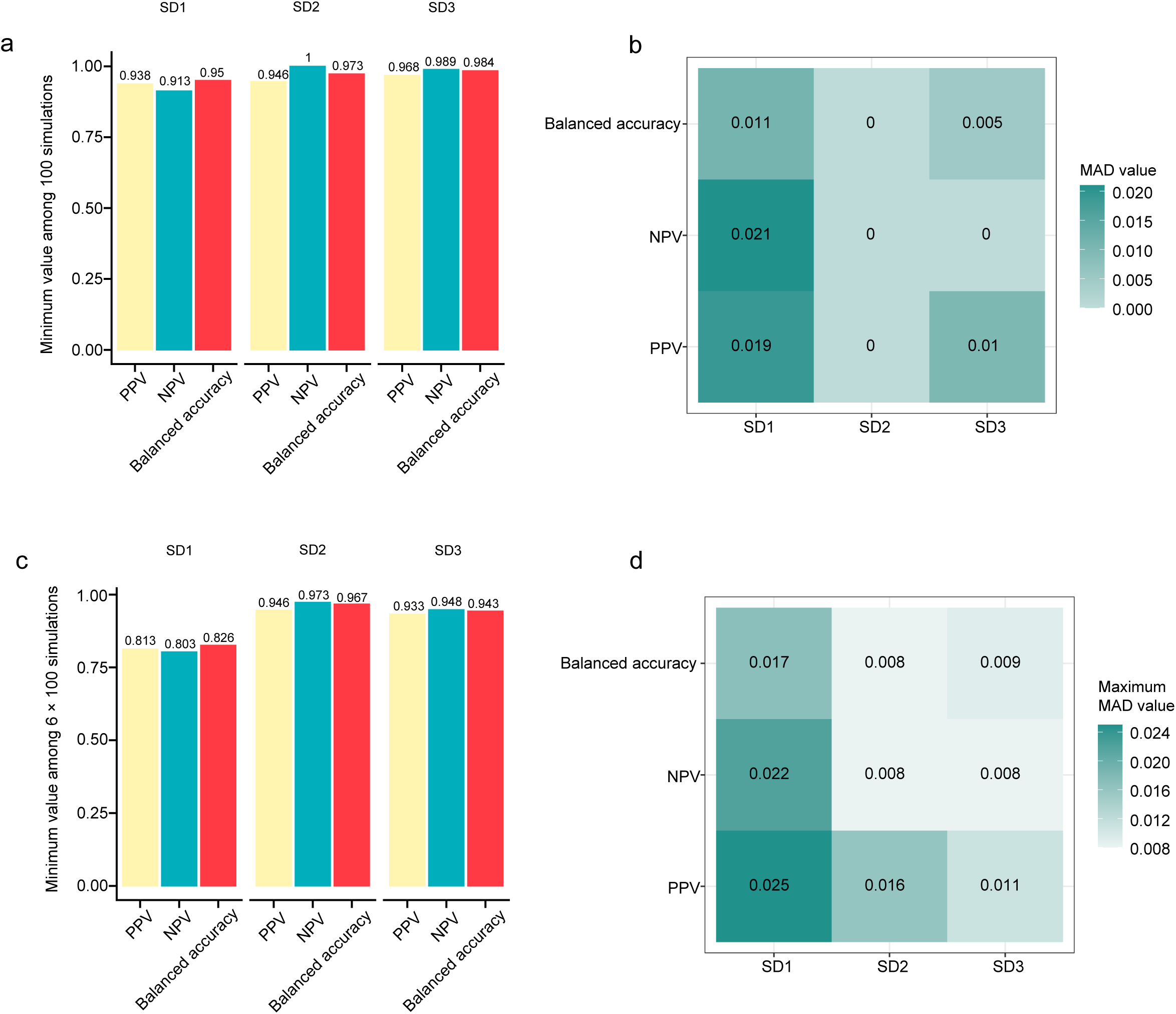
ScPP shows high prediction accuracy and stability in simulated datasets. **a,** Bar plots showing the minimum PPV, NPV and balanced accuracy among 100 simulations with default parameter α = 0.2 in SD1, SD2 and SD3. **b,** Heatmap showing the MAD of PPV, NPV and balanced accuracy among 100 simulations with default parameter α = 0.2 in SD1, SD2 and SD3. **c,** Bar plots showing the minimum PPV, NPV and balanced accuracy among 6×100 simulations under six different α values (0.2, 0.25, 0.3, 0.35, 0.4 and 0.45) in SD1, SD2 and SD3. **d,** Heatmap showing the maximum MAD of PPV, NPV and balanced accuracy among 6×100 simulations under the six α values in SD1, SD2 and SD3. MAD, median absolute deviation. PPV, positive predictive value. NPV, negative predictive value.

Next, we tested the performance of ScPP under different α values in the simulated datasets. We set the α value as 0.2, 0.25, 0.3, 0.35, 0.4 and 0.45, respectively. Results showed that the minimum PPV was 0.813, 0.946 and 0.933 in SD1, SD2 and SD3, respectively, among 6 × 100 simulations; the minimum NPV was 0.803, 0.973 and 0.948, respectively; and the minimum balanced accuracy was 0.826, 0.967 and 0.943 respectively (Fig. 2c). Likewise, the results indicated the high prediction stability of ScPP. The maximum MAD of PPV for these different α values was 0.025, 0.016 and 0.011 in SD1, SD2 and SD3, respectively; the maximum MAD of NPV was 0.022, 0.008 and 0.008, respectively; and the maximum MAD of balanced accuracy was 0.017, 0.008 and 0.009, respectively (Fig. 2d).

Taken together, these results demonstrate the high accuracy and stability of ScPP in predicting single cells’ phenotypes.

### Predicting malignant and non-malignant cells by ScPP

In real datasets, we first tested the ability of ScPP to identify malignant and non-malignant cells using a LUAD scRNA-seq dataset^32^. This dataset contained 34,063 LUAD and non-LUAD cells^48^. Based on the TCGA-LUAD bulk RNA-seq dataset, we identified two sets of marker genes for LUAD and non-LUAD, respectively. Using the marker gene sets, ScPP predicted 4,786 cells as LUAD cells (LUAD+) and 3,412 as non-LUAD cells (LUAD-) (Fig. 3a). Based on the cell type identification data provided in the original publication^32^, 83.41% of LUAD+ cells were verified to be cancer cells, and 100% of LUAD-cells to be non-cancer cells (Fig. 3b). The LUAD-cells harbored multiple cell types, including myeloid cells (66.2%), endothelial cells (17.8%), alveolar cells (10%), fibroblasts (3.6%) and T cells (1.7%) (Fig. 3b). Hypergeometric test indicated the results predicted by ScPP not to be achieved by chance (*P* < 2.2 × 10^−16^).

**Fig. 3.**
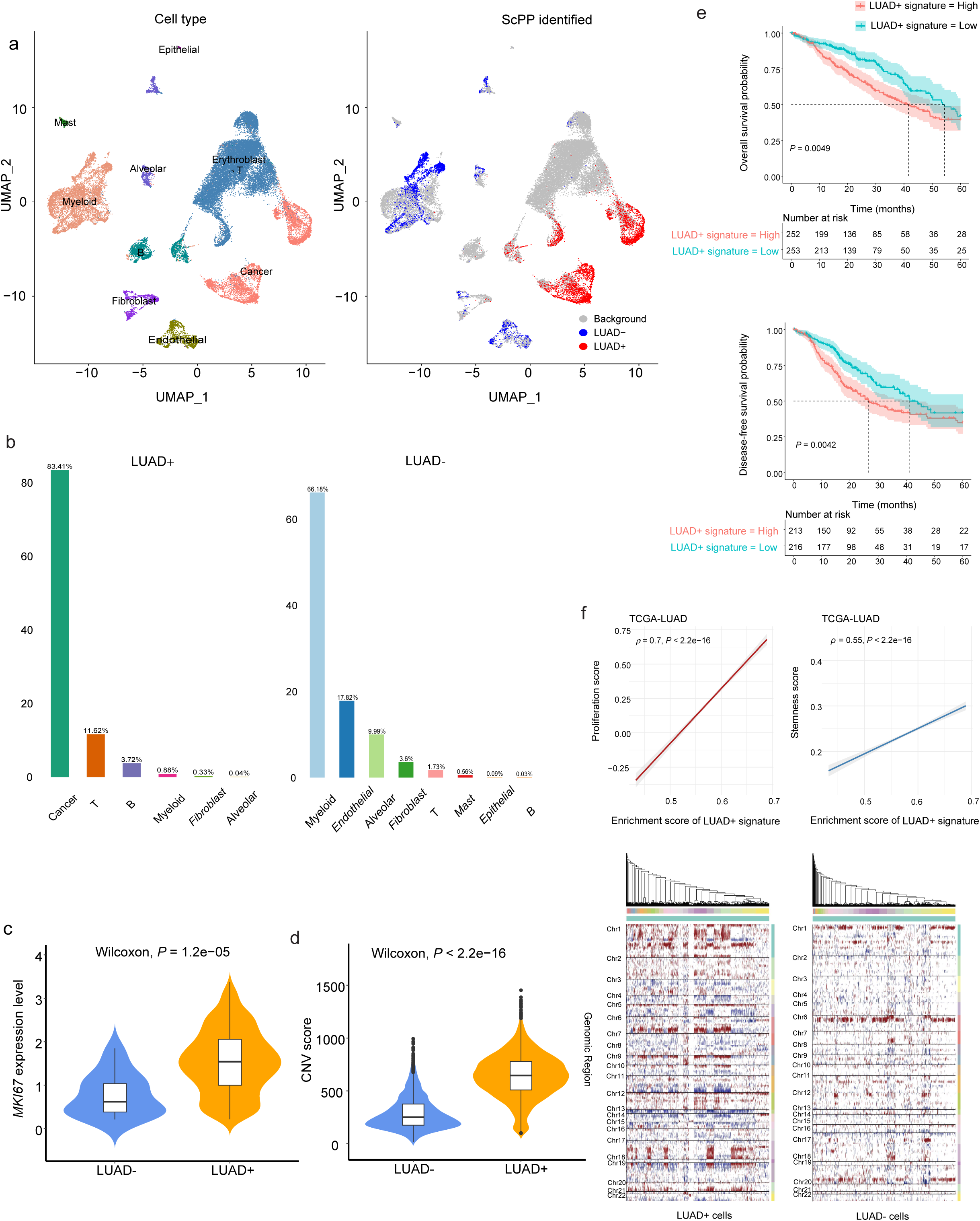
Predicting malignant and non-malignant cells in LUAD by ScPP. **a,** Uniform Manifold Approximation and Projection (UMAP) visualization of cells from LUAD scRNA-seq (left), and the cell subpopulations identified by ScPP (right). **b,** Bar plots showing the compositions of LUAD+ (left) and LUAD-(right) cells, with the percentage of each cell type over total LUAD+ or LUAD-cells indicating on the top of the bar. **c,** Violin plot showing the significantly higher *MKI67* expression levels in LUAD+ than in LUAD-cells. The one-tailed Mann-Whitney *U* test *P* value is shown. **d,** LUAD+ cells have greater CNVs than LUAD-cells. The one-tailed Mann-Whitney *U* test *P* value is shown. **e,** Kaplan-Meier curves displaying that tumors with higher enrichment scores (> median) of LUAD+ signature have worse 5-year survival prognosis than those with lower enrichment scores (< median). The log-rank test *P* value is shown. **f,** The enrichment scores of LUAD+ signature showing positive correlations with proliferation scores and stemness scores. The Spearman correlation coefficients (ρ) and *P* values are shown.

To further assess the facticity of ScPP-identified cells, we compared the expression levels of *MKI67*, a proliferation marker gene, between LUAD+ and LUAD-cells. This analysis showed that LUAD+ cells had significantly higher expression levels of *MKI67* (one-tailed Mann-Whitney *U* test, *P* = 1.2 × 10^−5^) (Fig. 3c). Moreover, the inferred CNVs by inferCNV^46^ were significantly higher in LUAD+ than in LUAD-cells (one-tailed Mann-Whitney *U* test, *P* < 2.2 × 10^−16^) (Fig. 3d). These results proved the prominent malignant property of LUAD+ cells. Distinct from bulk transcriptomes which are mixtures of the gene expression profiling in different cell types (such as tumor cells, immune cells and stromal cells), single-cell transcriptomes are the gene expression profiling in individual cells and thus more precisely reflect gene expression features of specific cell types. Therefore, we compared the gene expression profiling between LUAD+ and LUAD-cells to identify their marker genes. This analysis detected 183 upregulated and 270 downregulated genes in LUAD+ versus LUAD-cells (one-tailed Mann-Whitney *U* test, FDR < 0.05). As expected, many of the upregulated genes were adverse prognostic factors in TCGA-LUAD, and many of the downregulated genes were positive prognostic factors (Supplementary Fig. S1). Furthermore, the gene set enrichment scores of the 183 upregulated genes showed a negative correlation with survival prognosis (log-rank test, *P* = 0.0049 and 0.0042 for 5-year overall and disease-free survival, respectively) (Fig. 3e) but positive correlations with the proliferation scores (Spearman correlation ρ = 0.7; *P* < 2.2 × 10^−16^) and stemness scores (ρ = 0.55; *P* < 2.2 × 10^−16^) in TCGA-LUAD (Fig. 3f). The proliferation and stemness scores were the ssGSEA scores of their marker genes (Supplementary Table S3). These results collectively suggest that the LUAD+ and LUAD-cells predicted by ScPP are endowed with malignant and non-malignant properties, respectively.

### Identifying cell subpopulations associated with the ER status in breast cancer by ScPP

Breast cancer (BC) is the most common malignancy in female and also the most common cancer overall, accounting for 31% of cancer diagnoses^49^. Different subtypes of BC have markedly different prognoses and therapeutic regimens^50^. Here we used ScPP to identify ER-positive (ER+) and ER-negative (ER-) BC cells in the scRNA-seq dataset GSE176078^35^. This dataset contained gene expression profiles of eight major cell types, including cancer epithelial cells, endothelial cells, myeloid cells, plasmablasts, T cells, B cells, cancer-associated fibroblasts (CAFs) and mesenchymal cells^35^. We analyzed the 22,025 cancer epithelial cells to identify their ER status-associated phenotype (Fig. 4a). Likewise, ScPP first yielded ER+ and ER-marker genes by analyzing the TCGA BC (TCGA-BRCA) bulk RNA-seq dataset, which contained 1,050 BC samples. Based on the enrichment scores of these marker genes, ScPP predicted 2,598 cells to be ER+ and 3,135 cells ER-(Fig. 4b). As anticipated, more than 97% of the ER+ cells were verified to be luminal A or luminal B cells, and more than 98% ER-cells were basal-like or cycling cells (Fig. 4c). Notably, all the ER+ cells were derived from ER+ BCs, and almost all (> 99.9%) of ER-cells were from triple-negative BCs (Fig. 4d). Again, hypergeometric test suggested that these results predicted by ScPP were not likely to be gained by chance (*P* < 2.2 × 10^−16^).

**Fig. 4.**
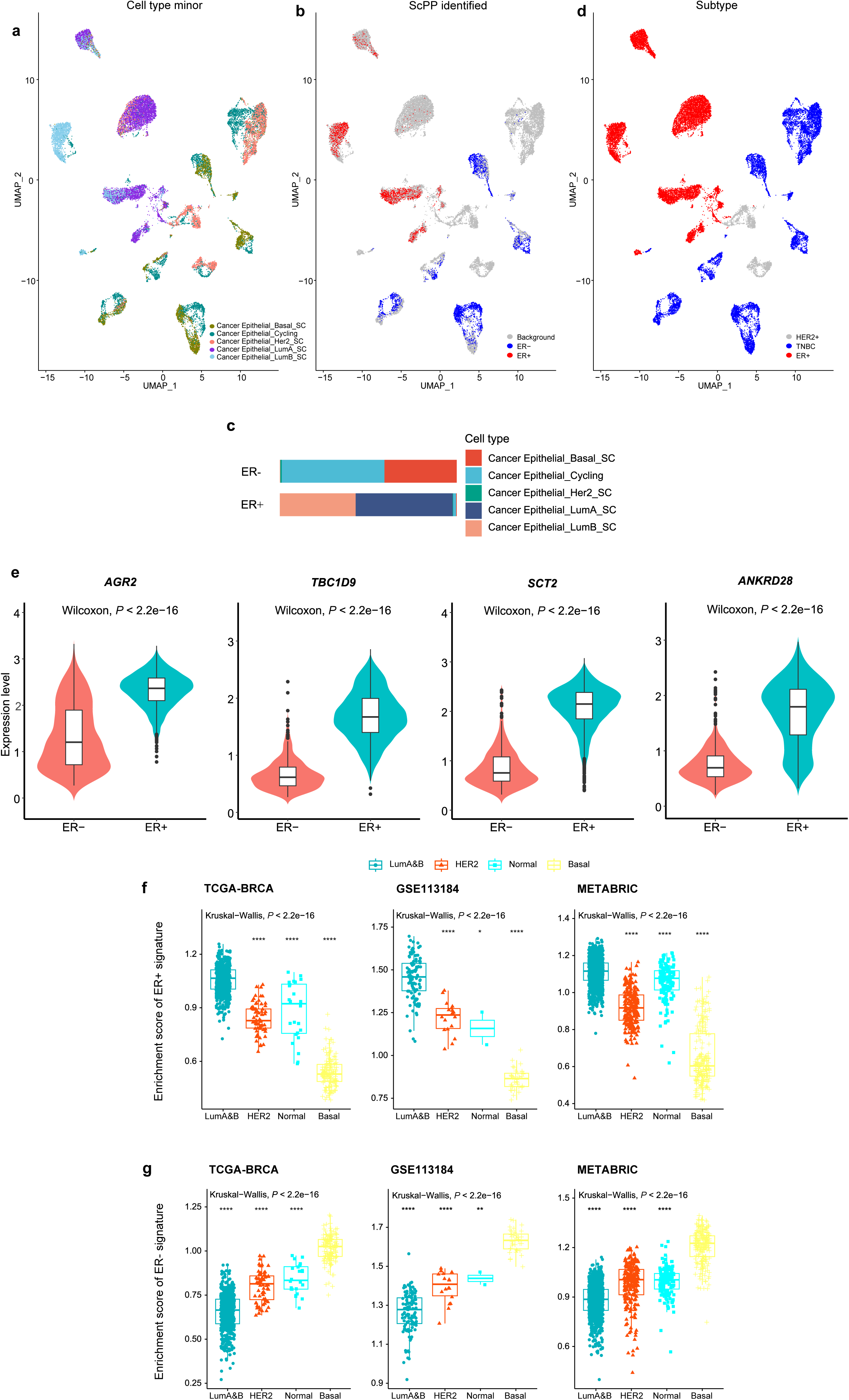
Identifying cell subpopulations associated with the ER status in breast cancer by ScPP. **a,** UMAP visualization of 22,025 cancer epithelial cells with different minor cell types from the BRCA scRNA-seq. **b,** UMAP visualization of ScPP-identified cell subpopulations associated with the ER status. **c,** Cell compositions of ScPP-identified ER+ and ER-cells. **d,** UMAP showing that a majority of ER+ cells identified by ScPP are from ER+ patients, and most of ER-cells are from TNBC patients. **e,** Violin plots to compare the expression levels of several upregulated genes in ER+ cells between ER+ and ER-cells. The one-tailed Mann-Whitney *U* test *P* values are shown. Comparisons of the enrichment scores of ER+ signature (**f**) and ER-signature (**g**) among the breast cancer subtypes. The Kruskal-Wallis test *P* values (**f, g**) and one-tailed Mann-Whitney *U* test *P* values for comparing luminal A&B (**f**) or basal (**g**) with other subtypes are shown. * *P* < 0.05, ** *P* < 0.01, *** *P* < 0.001, *** *P* < 0.0001. It also applies to the following figures.

To evaluate the transcriptional differences between the ER+ and ER-cells identified by ScPP, we compared gene expressions between both groups of cells. This analysis identified 239 upregulated and 171 downregulated genes in ER+ versus ER-cells (one-tailed Mann-Whitney *U* test, FDR < 0.05). The 239 upregulated genes in ER+ cells included several genes reported to be important in ER+ BCs, such as *AGR2*^51^, *TBC1D9*^52^, *STC2*^53,54^ and *ANKRD28*^55^ (Fig. 4e). Furthermore, in three independent BC bulk datasets (TCGA-BRCA, METABRIC^23^ and GSE113184^24^), the enrichment levels of ER+ cell signatures, namely the 239 genes upregulated in ER+ cells, were significantly higher in luminal A or luminal B than in other subtypes (one-tailed Mann-Whitney *U* test, *P* < 2.2 × 10^−16^) (Fig. 4f). In contrast, the ER-cell signatures, namely the 171 genes downregulated in the ER+ cells, showed significantly higher enrichment in basal-like than in other subtypes (*P* < 2.2 × 10^−16^) (Fig. 4g). Taken together, these data demonstrate that the identification of ER-associated cell subpopulations by ScPP is honest.

### Identifying cell subpopulations associated with the MSI status in colorectal cancer by ScPP

MSI is a typical property of the cancers with mismatch-repair deficiency and a useful biomarker for cancer management in multiple cancer types, including colorectal, endometrial, and gastric cancers^56,57^. We employed ScPP to identify cell subpopulations associated with the MSI status in the scRNA-seq dataset GSE132465 for CRC^37^. Similarly, ScPP first yielded the marker genes of MSI-high (MSI+) and MSI-low/microsatellite stable (MSI-) tumors by analyzing the TCGA CRC bulk RNA-seq dataset, which contained 377 CRC samples. Based on the enrichment scores of the marker genes, among 17,469 malignant epithelial cells in GSE132465 (Fig. 5a), ScPP identified 1,622 MSI+ cells and 1,615 MSI-cells (Fig. 5b). We compared the gene expression profiles between the MSI+ and MSI-cells, and found 93 and 29 genes significantly upregulated in MSI+ and MSI-cells, respectively. We defined the 93 and 29 genes as MSI+ and MSI-signatures, respectively. Notably, some of the MSI+ signatures have been reported to be overexpressed in MSI-high tumors, such as *AGR2*^58^, *PLAC8*^59^ and *REG4*^60^ (Fig. 5c). As expected, the enrichment scores of MSI+ signatures were noticeably higher in MSI-high than in MSI-low/microsatellite stable CRC patients in TCGA-CRC, while MSI-signatures exhibited an opposite pattern (one-tailed Mann-Whitney *U* test, *P* < 2.2 × 10^−16^) (Fig. 5d). Moreover, MSI+ signatures displayed significant negative correlations of their enrichment scores with the expression levels of three mismatch repair genes in TCGA-CRC, including *MSH2*, *PMS2* and *MLH1* (Spearman correlation, *P* < 0.05) (Fig. 5e). Furthermore, we observed a significant positive correlation between the enrichment scores of MSI+ signatures and TMB in TCGA-CRC (Spearman correlation ρ = 0.38; *P* = 1.4 × 10^−11^) (Fig. 5f), conforming to the fact that MSI-high tumors have increased TMB^61^. Also, MSI is a reflection of genomic instability^62^ which in turn may result in high ITH. Expectedly, we found a significant positive correlation between the MSI+ signatures’ enrichment and ITH scores in TCGA-CRC (ρ = 0.17; *P* = 0.0007) (Fig. 5g). Taken together, these results suggest that the MSI+ CRC cells predicted by ScPP have the primary properties of MSI tumors.

**Fig. 5.**
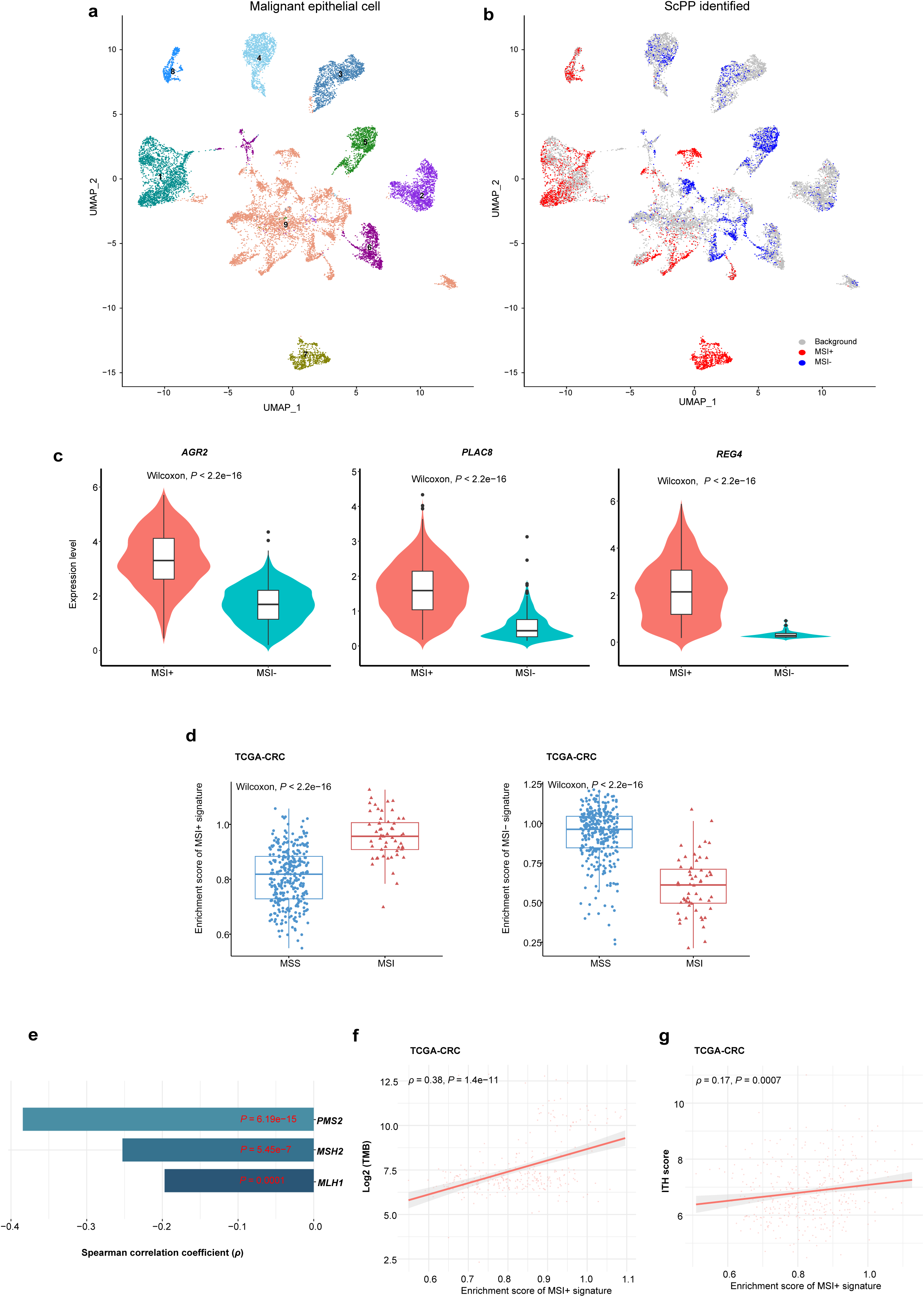
Identifying cell subpopulations associated with the MSI status in colorectal cancer by ScPP. **a,** UMAP visualization of 17,469 heterogeneous epithelial cells from CRC. **b,** UMAP visualization of ScPP-identified cell subpopulations associated with the MSI status. **c,** Comparisons of the expression levels of several upregulated genes in MSI+ cells between MSI+ and MSI-cells. **d,** Comparisons of the enrichment scores of MSI+ and MSI-signatures between MSI-high and MSS CRC patients. **e-g,** Spearman correlations between the enrichment scores of MSI+ signatures and the expression levels of three DNA mismatch repair genes (**e**), TMB (**f**), and ITH scores (**g**) in TCGA-CRC. The one-tailed Mann-Whitney *U* test *P* values are shown in (**c, d**), and the Spearman correlation coefficients (ρ) and *P* values are shown in (**e-g**). MSI: microsatellite instability. MSS: microsatellite stable. TMB: tumor mutation burden. ITH: intratumor heterogeneity.

### Identifying cell subpopulations associated with CNV in melanoma by ScPP

CNV reflects genomic instability that is a prominent property of most cancers^63^. We applied ScPP to a scRNA-seq dataset GSE132465^36^ for melanoma which contained 2,018 tumor cells. We first identified the marker genes of CNV-high (CNV+) and CNV-low (CNV-) tumors using the TCGA SKCM bulk RNA-seq dataset containing 211 SKCM samples by Spearman correlation analysis (FDR < 0.05; |ρ| > 0.2). Based on the enrichment scores of the marker genes, ScPP identified 284 CNV+ and 200 CNV-tumor cells in GSE132465 (Fig. 6a). Expectedly, the inferred CNVs by inferCNV^46^ were significantly higher in CNV+ than in CNV-cells (*P* < 2.2 × 10^−16^) (Fig. 6b). Likewise, we identified 259 CNV+ signatures and 229 CNV-signatures by a gene differential expression analysis of the CNV+ and CNV-cells. We found the enrichment scores of CNV+ signatures to correlate negatively with the enrichment scores of antitumor immune signatures (CD8+ T cells, cytolytic activity, human leukocyte antigen (HLA), and interferon (IFN) response) in TCGA-SKCM (ρ < −0.2; *P* < 0.0001), while the CNV-signatures correlated positively with the antitumor immune signatures (ρ > 0.5; *P* < 2.2 × 10^−16^) (Fig. 6c). These results support that increased CNV is associated with reduced antitumor immune responses^64^. In addition, the enrichment scores of CNV+ signatures were significantly lower in melanoma patients responsive to ICIs than in those unresponsive to ICIs (one-tailed Mann-Whitney *U* test, *P* = 0.03), while the CNV-signatures showed a contrary result (*P* = 0.04) (Fig. 6d). Again, it supports that increased CNV is associated with reduced immunotherapy responses^64^. Furthermore, the enrichment scores of CNV+ signatures correlated positively with the scores of homologous recombination deficiency (HRD) (ρ = 0.18; *P* = 1.4 × 10^−4^) and ITH in TCGA-SKCM (ρ = 0.25; *P* = 7.2 × 10^−8^), in contrast to the CNV-signatures correlating negatively with them (HRD: ρ = −0.09 and *P* = 0.04; ITH: ρ = −0.41 and *P* < 2.2 × 10^−16^) (Fig. 6e). Since HRD is a major cause of large-scale genomic instability^65^ and ITH is a consequence of genomic instability^66^, these results supported the positive association between CNV and genomic instability. Overall, these results suggest that the CNV-associated cell subpopulations inferred by ScPP are characterized by CNV-related properties in cancer.

**Fig. 6.**
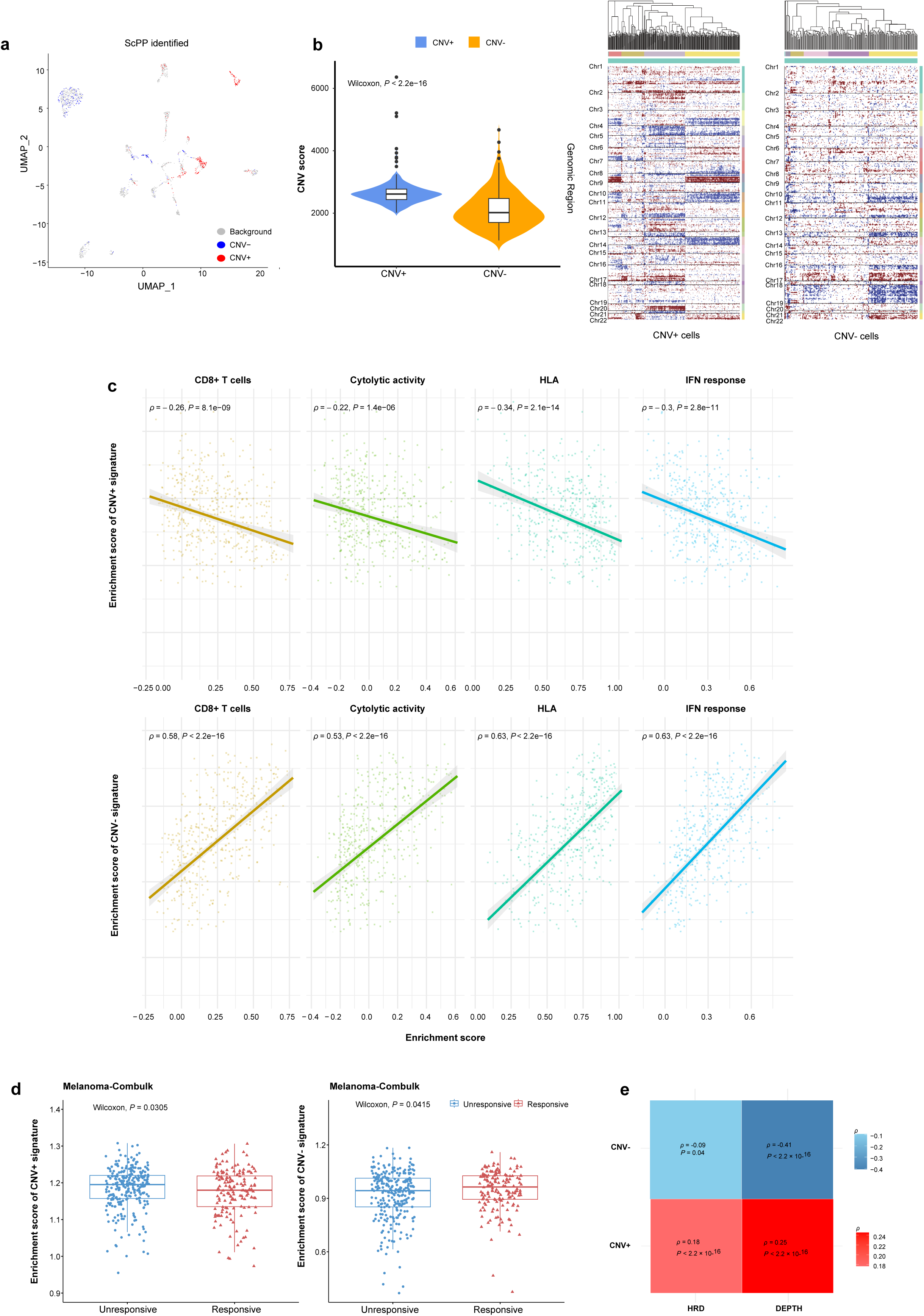
Identifying cell subpopulations associated with CNV in melanoma by ScPP. **a,** UMAP visualization of ScPP-identified cell subpopulations associated with ScPP-identified cell subpopulations associated with CNV from 2,018 melanoma cells. **b,** The inferred CNV scores are significantly higher in CNV+ than in CNV-cells. **c,** The enrichment scores of antitumor immune signatures correlate negatively with CNV+ signatures and correlate positively with CNV-signatures. **d,** Compared to the melanoma patients not responding to ICIs, those responding to ICIs show lower enrichment of CNV+ signatures but higher enrichment of CNV-signatures. **e,** Both HRD and ITH correlates positively with the enrichment of CNV+ signatures but negatively with the CNV-signatures. The one-tailed Mann-Whitney *U* test *P* values are shown in (**b, d**), and the Spearman correlation coefficients (ρ) and *P* values are shown in (**c, e**).

### Identifying cell subpopulations related to survival prognosis in LUAD by ScPP

To identify survival-related cell subpopulations in the LUAD scRNA-seq dataset^48^ analyzed previously, ScPP first analyzed the TCGA-LUAD bulk dataset to identify the marker genes whose expression was significantly associated with overall survival using univariate Cox regression analysis. This analysis detected 446 and 230 marker genes which were positive and adverse prognostic factors in LUAD, respectively. Based on the enrichment scores of the marker genes, ScPP predicted 1,226 cells related to worse survival and 1,090 cells associated with better survival in the 34,063 cells (Fig. 7a). Here we termed the group of cells associated with worse survival LUAD-WS and the group of cells associated with better survival LUAD-BS. Of the 1,226 cells in LUAD-WS, 635 (51.8%) were malignant cells, 310 (25.3%) were T cells, 225 (18.4%) were myeloid cells, 31 (2.5%) were endothelial cells and 23 (1.9%) were fibroblasts (Fig. 7b). This observation suggests that both tumor cells and non-tumor cells in the tumor microenvironment may contribute to cancer patients’ prognosis. Of the 1,090 cells in LUAD-BS, more than 70% were immune cells, including 450 (41.3%) T cells, 299 (27.4%) B cells and 41 (3.8%) mast cells (Fig. 7b). It indicates that active antitumor immune responses may lead to a better prognosis in LUAD. Interestingly, many B cells occurred in LUAD-BS but only two B cells included in LUAD-WS. It is consistent with a previous report that the density of T cells surrounded by B-cell follicles is associated with favorable clinical outcomes in LUAD^67^.

**Fig. 7.**
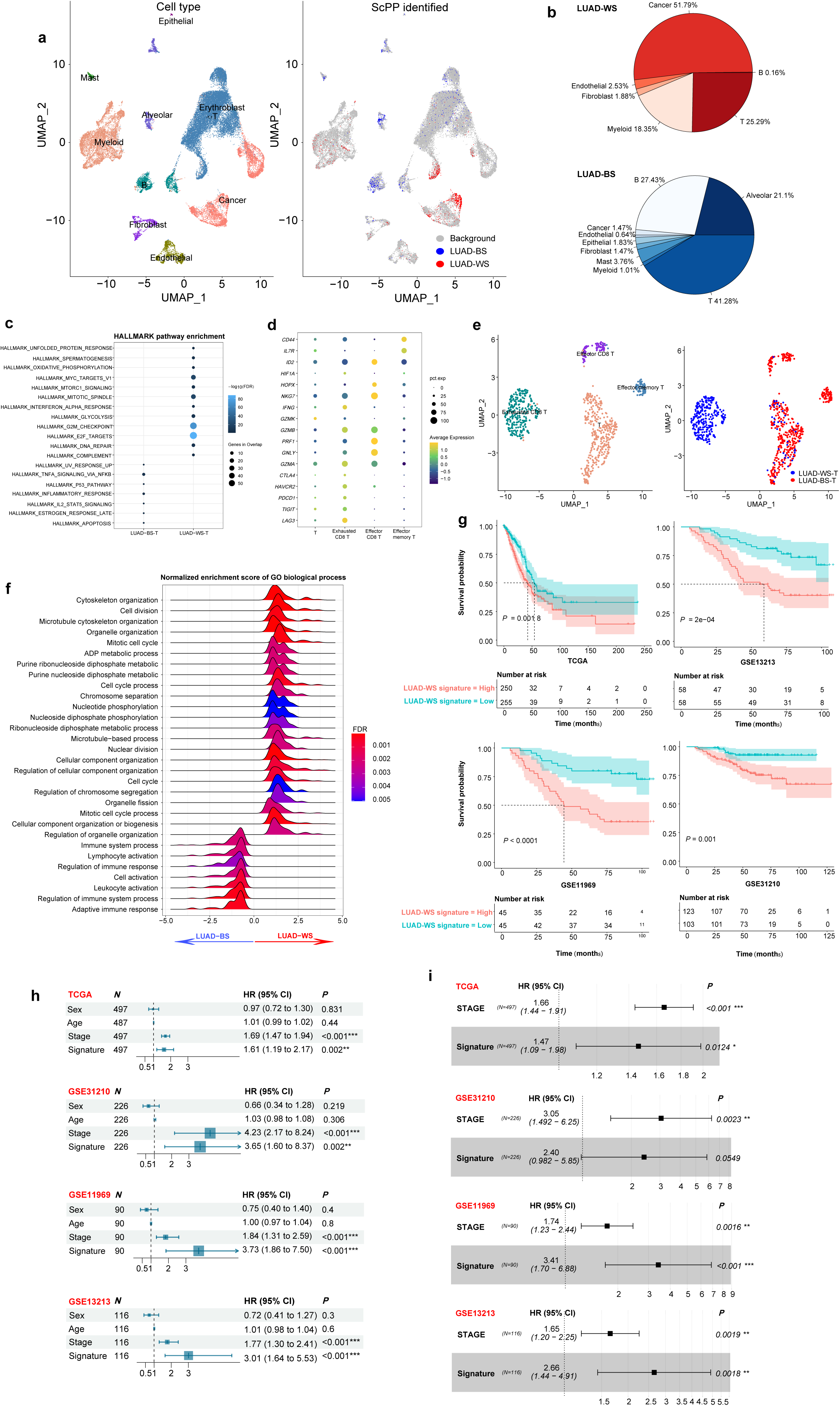
Identifying cell subpopulations associated with survival prognosis in LUAD by ScPP. **a,** UMAP visualization of cells from LUAD scRNA-seq (left), and the ScPP-identified survival-related cells (right). **b,** Pie plots showing the compositions of the group of cells associated with worse survival (LUAD-WS cells, up) and the group of cells associated with better survival (LUAD-BS cells, down); the percentage of each cell type over LUAD-WS or LUAD-BS cells are indicated beside the pie. **c,** Dot plot showing the hallmark pathways enriched in LUAD-WS and LUAD-BS T cells. **d,** Dot plot showing the expression patterns of the gene markers of T cell subtypes. **e,** UMAP visualization of T cell subtypes (left), and ScPP-identified survival-related T cells (right). **f,** Ridge plot showing the GO biological processes enriched in LUAD-WS and LUAD-BS cells. **g,** Kaplan-Meier curves to compare overall survival (OS) time between LUAD patients with high enrichment scores of LUAD-WS marker genes (> median) and those with low enrichment scores (< median) in four independent LUAD datasets. The log-rank test *P* values are shown. **h,** Univariate Cox proportional hazards regression analysis with the response variable “OS time” and the predictor variables “sex,” “age,” “stage” and “signature.” sex: “Male” = 1 and “Female” = 2; age: the age at diagonosis; stage: the pathological stage (Stage I = 1, Stage II = 2, Stage III = 3, and Stage IV = 4); signature: the ssGSEA scores of LUAD-WS marker genes. **i,** Multivariable Cox proportional hazards regression analysis with the response variable “OS time” and the predictor variables “stage” and “signature.” The two-tailed Wald test *P* values are shown. HR: hazard ratio. CI: confidence interval.

Intriguingly, we found that both groups of survival-related cells included a fraction of T cells (310 in LUAD-WS and 450 in LUAD-BS). To explore the difference in functions between both groups of T cells, we compared their gene expression prefiles, and detected 136 and 25 genes significantly upregulated in LUAD-WS and LUAD-BS T cells, respectively. Gene set enrichment analysis by GSEA^68^ showed 12 pathways having significant correlations with the 136 genes upregulated in LUAD-WS T cells (FDR < 0.05) (Fig. 7c). These pathways were mainly related to cell cycle, DNA repair, and glycolysis. In contrast, the 25 genes upregulated in LUAD-BS T cells were enriched in seven pathways primarily involved in p53 signaling and immune/inflammatory responses (Fig. 7c). To further explore the survival-related T cells, we performed a cluster analysis of the 760 ScPP-identified T cells. This analysis identified four clusters of the T cells, namely exhausted CD8 T cells, effector CD8 T cells, effector memory T cells and other T cells (Fig. 7d-e), based on the expressions of related marker genes^44^. Remarkably, a majority (83.6%) of LUAD-WS T cells were exhausted T cells, while LUAD-BS T cells did not harbor any exhausted T cells. On the other hand, a minority (0.32%) of LUAD-WS T cells were effector T cells, compared to 16% of LUAD-BS T cells being effector T cells (Fig. 7e). These results are reasonable since activated cytotoxic T cells play a crucial role in antitumor immune responses, while exhausted T cells have a role in antitumor immunosuppression^69^. Thus, this analysis indicates that ScPP can effectively recognize cell subpopulations related to cancer prognosis.

We further compared the gene expression profiles between LUAD-WS and LUAD-BS. This analysis detected 153 genes upregulated in LUAD-WS (termed LUAD-WS marker genes) and 56 genes upregulated in LUAD-BS (termed LUAD-BS marker genes) (one-tailed Mann-Whitney *U* test, FDR < 0.05). Functional analysis of these marker genes showed that the cell cycle, cell devision and cell growth regulation were highly activated in LUAD-WS, while immune-related biological processes were highly activated in LUAD-BS, such as adaptive immune response, lymphocyte activation and immune system process (Fig. 7f). Furthermore, in four independent LUAD bulk datasets, the enrichment scores of LUAD-WS marker genes displayed significant inverse correlations with overall survival time (log-rank test, *P* < 0.05) (Fig. 7g). In contrast, the enrichment scores of LUAD-BS marker genes showed significant positive correlations with overall survival in two of the four datasets (*P* < 0.05) (Supplementary Fig. S2). To compare the survival relevance between the LUAD-WS markers and other clinical parameters, we used univariate Cox survival analyses to assess the prognositic value of sex, age, pathological stage, and LUAD-WS markers in the four datasets. This analysis revealed that only pathological stage and LUAD-WS markers had significant correlations with survival (Fig. 7h). Furthermore, multivariable Cox survival analyses showed that LUAD-WS markers remained an adverse prognostic factor after adjusting for pathological stage in three of the four datasets (Fig. 7i). Taken together, these results suggest that the survival-related cell subpopulations inferred by ScPP have authentic prognostic relevance.

### Identifying cell subpopulations related to immunotherapy response in melanoma by ScPP

Immunotherapy, such as immune checkpoint blockade^70^, has achieved success in treating a varity of cancers. However, a large proportion of cancer patients fail to respond to ICIs. Hence, identifying predictive markers for immunotherapy response is crucial. Here we tested ScPP in recognizing single cells related to immunotherapy response in the scRNA-seq dataset GSE115978^36^ for melanoma. The reference bulk dataset was the gene expression profiles in 427 melanoma patients treated with ICIs, including 158 responders and 269 non-responders. This dataset was generated by combining six bulk RNA-seq datasets and correcting for batch effects (see Methods). Likewise, we identified marker genes for responsive and unresponsive groups, respectively. Based on these markers, ScPP identified 1,407 responsive cells and 1,440 unresponsive cells from 7,186 cells. Notably, most of the responsive cells were immune cells, including CD8+ T cells (47.8%), B cells (27.6%), CD4+ T cells (8.4%), T cells (6.8%), macrophages (6.7%) and NK cells (1.6%) (Fig. 8a). In sharp contrast, most of the unresponsive cells were malignant cells (90.8%), and some were CAFs (4.2%) and endothelial cells (0.8%) (Fig. 8a). These data support previous findings that tumor cells, CAFs and endothelial cells play important roles in tumor immunosuppression^71–76^. Furthermore, cell-cell communication network analysis showed the responsive cells to have higher interaction weights with various immune cells and the unresponsive cells to interact more actively with CAFs and endothelial cells (Fig. 8b). Moreover, we analyzed changes in the signaling pattern of different cell types. This analysis found that some immune-stimulatory molecules were more highly activated in the responsive cells, such as MHC-I, MHC-II, IL16, CD45 and CD48, while immune-inhibitory molecules were more active in the unresponsive cells, such as cell adhesion molecules and macrophage migration inhibitory factors (Fig. 8c).

**Fig. 8.**
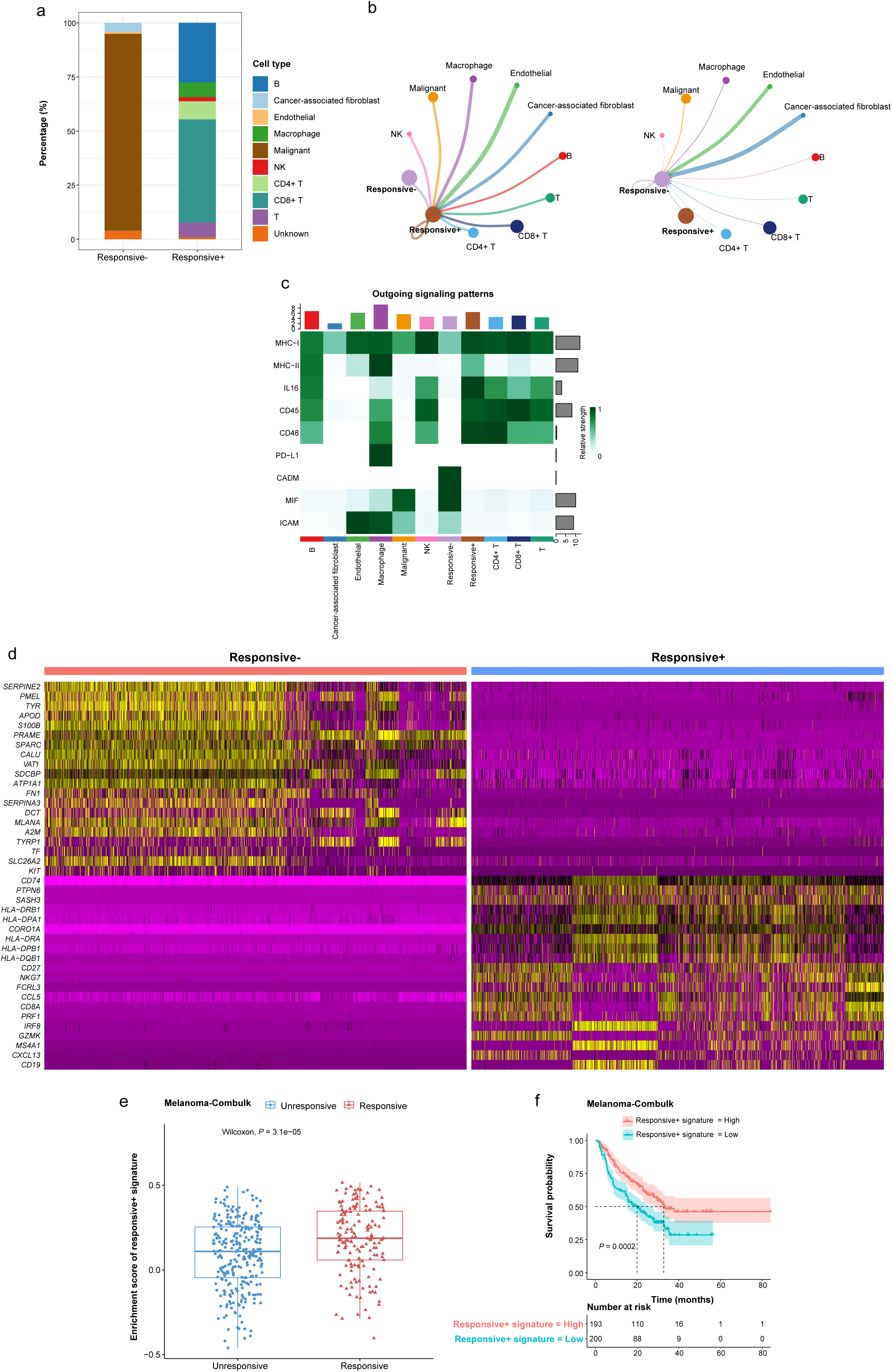
Identifying cell subpopulations associated with immunotherapy response in melanoma by ScPP. **a,** Cellular compositions of responsive+ and responsive-cells in melanoma identified by ScPP. **b,** Interaction weights between responsive+ or responsive-cells and other cells. The plots were generated by the R package “CellChat.” The line thickness indicates the interaction strength. **c,** Signaling patterns of responsive+ and responsive-cells. **d,** Heatmap showing the top 20 highly expressed genes in responsive+ and responsive-cells. **e,** Comparisons of the enrichment scores of responsive+ signatures between responsive and unresponsive patients in Melanoma-Combulk. The one-tailed Mann-Whitney *U* test *P* values are shown. **f,** Kaplan-Meier curves displaying that tumors with higher enrichment of responsive+ signatures have better survival prognosis than those with lower enrichment scores. The log-rank test *P* value is shown.

Differential gene expression analysis identified 443 and 275 genes to be upregulated in the responsive and unresponsive cells, respectively. We termed the 443 and 275 genes responsive+ and responsive-signatures, respectively. Notably, many of the responsive+ signatures have been reported to have an active role in immunotherapy response, such as *NKG7*^77^, *CCL5*^78^, *PRF1*^79^, *CD27*^80^ and *CD8A*^80^ (Fig. 8d). Furthermore, in the melanoma-Combulk cohort receiving the ICI treatment, the enrichment levels of responsive+ signatures were significantly higher in the responsive than in unresponsive patients (one-tailed Mann-Whitney *U* test, *P* = 3.1 × 10^−5^) (Fig. 8e). As a result, the patients with higher enrichment of responsive+ signatures displayed better survival (log-rank test, *P* = 0.0002) (Fig. 8f). Taken together, these results suggest that ScPP is capable of identifying cell subpopulations related to immunotherapy response in cancer.

### ScPP has advantages in identifying cell subpopulations related to specific phenotypes over other algorithms

We compared the performance of identifying phenotype-associated cell subpopulations among ScPP and other algorithms, including Scissor^13^ and scAB^14^, in both simulated and real datasets. First, we compared the performance of ScPP and Scissor in three simulated datasets, namely SD1, SD2 and SD3. Here we did not run scAB in simulated datasets considering that scAB only identifies phenotype+ but not phenotype-cell subpopulations. For both ScPP and Scissor, we analyzed the simulated datasets using six different while commonly used α values (0.2, 0.25, 0.3, 0.35, 0.4 and 0.45 for ScPP, and 0.005, 0.01, 0.05, 0.1, 0.2 and 0.3 for Scissor), respectively. Under each α value, both algorithms repeated running 100 times for the simulated datasets (see Methods). Thus, for each α value, each algorithm obtained 100 results related to each of the four simulated datasets. We calculated the average performance, namely average PPV, NPV and balanced accuracy, of the 100 results. In all cases, ScPP displayed significantly higher average performance than Scissor in the simulated datasets (one-tailed Mann-Whitney *U* test, *P* < 0.05) (Fig. 9a). Furthermore, in almost all cases, the MAD of the six median performance in 100 simulations under each of the six α values by ScPP was smaller than that by Scissor in the simulated datasets (Fig. 9b); and the MADs of the performances in 100 times of simulations under the same α values by ScPP were consistently smaller than those by Scissor in all three simulated datasets (Fig. 9c). These results collectively indicate that ScPP has higher accuracy and stability than Scissor in identifying phenotype-related cell subpopulations.

**Fig. 9.**
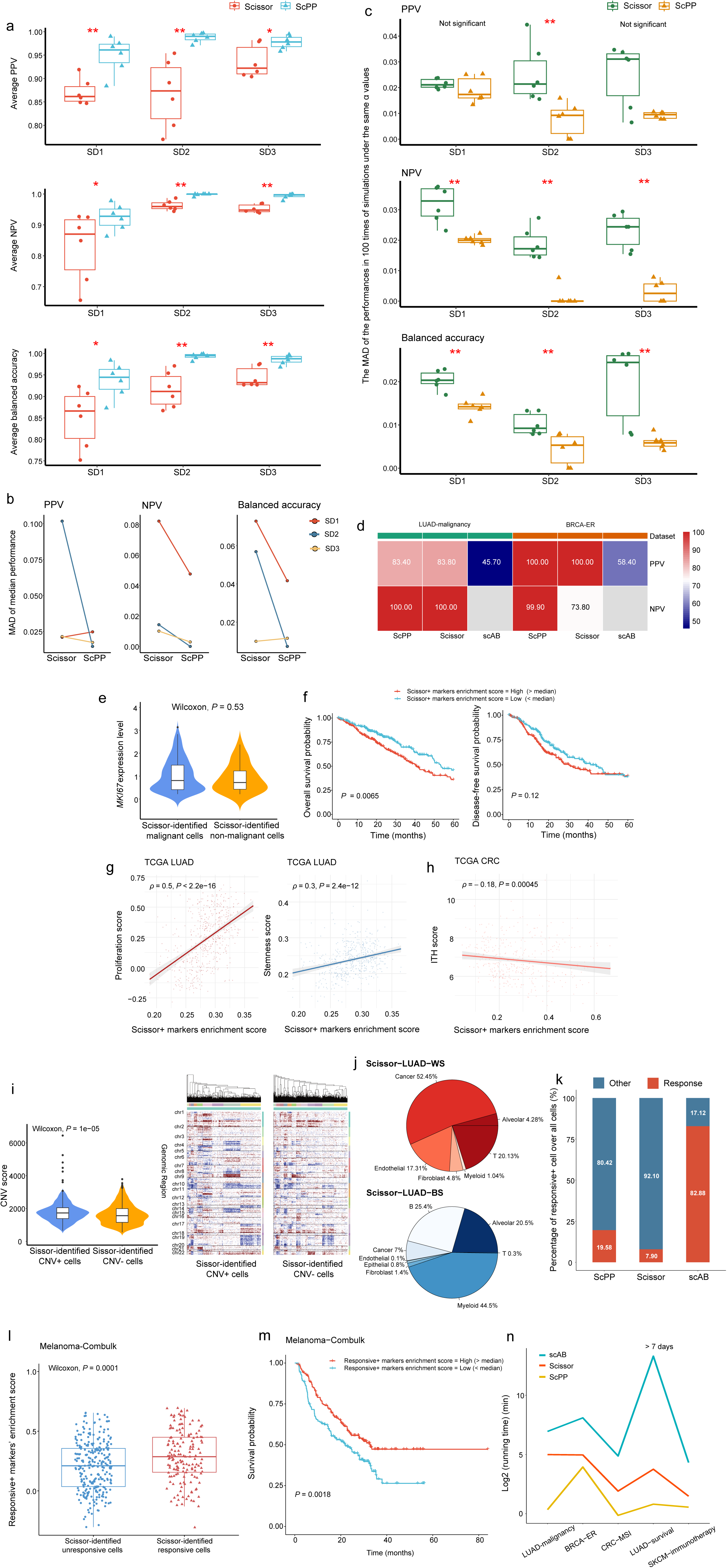
Comparisons between ScPP and two other algorithms. **a,** ScPP displays significantly higher average performance than Scissor in the simulated datasets. **b,** In 7 of 9 comparisons, the MAD of the six median performance in 100 simulations under each of the six α values by ScPP is smaller than that by Scissor in the simulated datasets. **c,** In almost all cases, the MADs of the performances in 100 times of simulations under the same α values by ScPP are significantly smaller than those by Scissor in three simulated datasets. **d,** ScPP showing the highest PPV and NPV in predicting cell subpopulations associated with malignancy and ER status among the three algorithms. **e,** The proliferation marker gene *MKI67* showing no significant expression difference between the malignant and non-maliganant cells identified by Scissor. **f,** Kaplan-Meier curves showing the correlation between the enrichment scores of the LUAD malignancy-associated Scissor+ markers and TCGA-LUAD patients’ survival prognosis. **g,** The correlation between the enrichment scores of the LUAD malignancy-associated Scissor+ markers and the proliferation and stemness scores. **h,** The negative correlation between the enrichment scores of CRC MSI-associated Scissor+ markers and ITH scores in TCGA-CRC. **i,** Comparison of inferCNV-infered CNV scores between CNV+ and CNV-cells identified by Scissor. **j,** Pie plots showing the compositions of the group of cells associated with worse survival (Scissor-LUAD-WS cells, up) and the group of cells associated with better survival (Scissor-LUAD-BS cells, down) identified by Scissor; the percentage of each cell type over Scissor-LUAD-WS or Scissor-LUAD-BS cells are indicated beside the pie. **k,** Percentages of responsive+ cells over all cells identified by the three algorithms, in which the prediction by ScPP is closest to the practical immunotherapy response rate in melanoma. **l-m,** The Scissor-identified responsive+ markers’ enrichment scores are significantly higher in the responsive than in unresponsive melanomas (**l**), and correlate positively with patients’ survival in melanoma-Combulk (**m**), but the corresponding *P* values are larger than those obtained in the ScPP-associated analyses. **n,** Comparion of running time of the algorithms in the same datasets. The one-tailed Mann-Whitney U test P values are shown in **(a, c, e, I, l)**, the log-rank test P value is shown in **(f, m)**, and the Spearman correlation coefficients (ρ) and *P* values are shown in **(g, h)**.

In the real datasets, we implemented Scissor and scAB to identify cell subpopulations related to malignancy, ER, MSI, CNV, survival, and immunotherapy response using the identical bulk and single-cell datasets as well as the same computational resources. We compared the results achieved by both algorithms with those by ScPP, as shown in the previous results. For Scissor, in each real dataset, we reported the results obtained under the optimal α value for the algorithm. The optimal α value was generated by Scissor by setting the parameters: cutoff = 0.2 and alpha = NULL, as recommended by the Scissor’s developers^13^. For scAB, we reported the results obtained under its default parameters. In predicting cell subpopulations related to malignancy in LUAD, Scissor identified 1,718 malignant and 2,264 non-malignant cells, compared to 4,786 malignant and 3,412 non-malignant cells by ScPP. In this dataset, both ScPP and Scissor achieved high PPVs and NPVs (PPV: 83.4% (ScPP) versus 83.8% (Scissor); NPV: 100% versus 100%) (Fig. 9d). However, the proliferation marker gene *MKI67* showed no significant expression difference between the malignant and non-maliganant cells identified by Scissor (Fig. 9e), as compared with that its expression levels were significantly higher in the malignant than in non-maliganant cells identified by ScPP (Fig. 3c). Furthermore, by comparing the gene expression profiles between the 1,718 malignant and 2,264 non-malignant cells identified by Scissor, we obtained the genes upregulated in malignant cells, termed Scissor+ markers. Although the enrichment scores of the Scissor+ markers also correlated negatively with TCGA-LUAD patients’ prognosis, the correlation was less significant than that between ScPP-identified markers and prognosis (overall survival: *P* = 0.0065 (Scissor) versus 0.0049 (ScPP); disease-free survival: *P* = 0.12 (Scissor) versus 0.0042 (ScPP)) (Fig. 9f & Fig. 3e). Meanwhile, although the enrichment scores of the Scissor+ markers correlated positively with the proliferation scores and stemness scores, the correlation coefficients were lower than those for ScPP-identified markers (Spearman correlation coefficients: 0.5 (Scissor) versus 0.7 (ScPP) for proliferation; 0.3 (Scissor) versus 0.55 (ScPP) for stemness) (Fig. 9g). These results indicate that ScPP is more accurate than Scissor in predicting cells associated with LUAD malignancy. In the BC scRNA-seq dataset (GSE176078), Scissor predicted 1,638 ER+ and 1,943 ER-cells, compared to 2,598 ER+ and 3,135 ER-cells predicted by ScPP; both ScPP and Scissor achieved 100% of PPV, while the NPV was lower with Scissor than with ScPP (73.8% versus 99.9%) (Fig. 9d). In predicting cell subpopulations related to MSI in CRC, Scissor identified 971 MSI+ and 2,045 MSI-cells, compared to 1,622 MSI+ and 1,615 MSI-cells by ScPP. By comparing the gene expression profiles between the MSI+ and MSI-cells identified by Scissor, we obtained the genes upregulated in the MSI+ cells, termed Scissor+ markers. Unexpectedly, the enrichment scores of Scissor+ markers correlated negatively with ITH scores in TCGA-CRC (Spearman correlation ρ = −0.18; *P* = 0.0005) (Fig. 9h), compared to a significant positive correlation between the MSI+ signatures’ enrichment and ITH scores (ρ = 0.17; *P* = 0.0007) (Fig. 5g). In predicting cell subpopulations related to CNV in CRC, Scissor identified 636 CNV+ and 681 CNV-cells, compared to 284 CNV+ and 200 CNV-cells by ScPP. Although the inferCNV-inferred CNV scores were higher in CNV+ than in CNV-cells identified by Scissor (*P* = 1 × 10^−5^), the difference was more significant between CNV+ and CNV-cells identified by ScPP (*P* < 2.2 × 10^−16^) (Fig. 9i & Fig. 6b). In predicting cell subpopulations related to survival in LUAD, Scissor identified 959 and 1,000 cells associated with worse and better survival, respectively (termed Scissor-LUAD-BS and Scissor-LUAD-WS cells, respectively), compared to 1,226 LUAD-BS and 1,090 LUAD-WS cells by ScPP. However, 20.1% of Scissor-LUAD-WS cells were T cells, compared to only 0.3% of the Scissor-LUAD-BS cells being T cells (Fig. 9j). In contrast, in the predictions by ScPP, the proportion of T cells in LUAD-WS and LUAD-BS was 25.3% and 41.3%, respectively (Fig. 7b). Evidently, the result predicted by ScPP is more reasonable than that by Scissor as the single cells associated with better survival should harbor a higher proportion of antitumor immune cells than those associated with worse survival. In predicting cell subpopulations related to immunotherapy response in melanoma, Scissor inferred 568 responsive and 703 unresponsive cells, compared to 1,407 responsive and 1,440 unresponsive cells inferred by ScPP (Fig. 9k). By comparing the gene expression profiles between the responsive and unresponsive cells predicted by Scissor, we obtained the genes upregulated in the responsive cells, termed responsive+ markers. While the responsive+ markers’ enrichment scores were significantly higher in the responsive than in unresponsive melanomas, and correlated positively with patients’ survival in melanoma-Combulk, the corresponding *P* values were larger than those obtained in the ScPP-associated analyses (Fig. 9l-m & Fig. 8e-f). Taken together, these comparisons suggest that ScPP is more powerful and accurate than Scissor in predicting phenotype-related cell subpopulations.

scAB predicts cell subpopulations with some phenotype but not those lack of the phenotype^14^. In the prediction of cell subpopulations related to six phenotypes (malignancy, ER, MSI, CNV, survival, and immunotherapy response), scAB only output results associated with malignancy, ER, MSI, and immunotherapy response; scAB could not predict CNV-associated cell subpopulations as CNV is a continuous variable; in addition, we could not obtain survival-related results after running scAB for a week. scAB predicted 3,819 LUAD malignant cells, 45.7% of which were verified to be malignant, compared to 83.4% of the 4,786 malignant cells predicted by ScPP being validated to be malignant (Fig. 9d). It suggested a substantially lower prediction precision of scAB versus ScPP in this respect. For the ER status, scAB predicted 3,317 ER+ BC cells, with a PPV of 58.4%, compared to 100% by ScPP (Fig. 9d). Furthermore, scAB predicted 749 MSI+ CRC cells, compared to 1,622 MSI+ cells predicted by ScPP. When comparing the gene expression profiles between the 749 MSI+ CRC cells inferred by scAB and other CRC cells, there was no upregulated gene detected in MSI+ cells using the FindMarkers function in the Seurat R package with the default of Mann-Whitney *U* test. In contrast, when comparing the gene expression profiles between the MSI+ and MSI-CRC cells predicted by ScPP but not by scAB, we obtained 92 and 28 genes significantly upregulated and downregulated in MSI+ cells, respectively, almost the same result as the previous analysis. This comparison suggested a higher predictive power of ScPP versus scAB in predicting MSI-associated cell subpopulations. Lastly, scAB predicted 5,956 immunotherapy responsive cells from a total of 7,186 cells, compared to 1,407 responsive cells by ScPP. That is, scAB inferred 82.9% cells to have a phenotype favoring immunotherapy response, compared to 20% inferred by ScPP (Fig. 9k). Evidently, our inference is closer to the practical immunotherapy response rate of 20-30% in melanoma^81^.

Impressively, compared to Scissor and scAB, ScPP needs much less computation time in identifying phenotype-associated cell subpopulations. In all six phenotypes-associated analyses of real datasets, ScPP spent less than 2 minutes (min) except the ER-associated prediction (15.38 min), while Scissor took 31.82 min, 31.18 min, 3.76 min, 13.51 min and 2.79 min to predict malignancy, ER, MSI, survival and immunotherapy response-associated cell subpopulations, respectively, and scAB took 123.34 min, 272.95 min, 29.32 min, > 7 days and 19.99 min to identify malignancy, ER, MSI, survival and immunotherapy response-associated cell subpopulations, respectively (Fig. 9n). These results suggest that ScPP is much more efficient than the established algorithms.

## Discussion

Here we developed the algorithm ScPP to identify phenotype-associated cell subpopulations by combining single-cell and bulk data. Distinct from established algorithms, ScPP infers single cells’ phenotypes based on the expression profiles of phenotype-associated marker genes in bulks and single cells. Our experiments demonstrate that ScPP can effectively recognize cell subpopulations with certain phenotypes, such as malignancy, ER status, MSI, CNV, survival prognosis and immunotherapy response. An outstanding merit of ScPP is its fast implementation, as evidenced by its significantly less running time than established algorithms, such as Scissor and scAB. This is important as the volume of scRNA-seq data is increasing exponentially. Besides, our experiments suggest that ScPP has more excellent predictive performance than the established algorithms, namely its higher predictive power, accuracy, stability and efficiency. In addition, ScPP can infer cell subpopulations related to the phenotypes characterized by continuous variables; that may result in its wider range of applications.

We argue that the superiority of ScPP is ascribed to its simple and clear algorithm logic to deal with complex and heterogeneous data. Unlike other algorithms, such as Scissor and scAB, ScPP does not use pairwise correlations to link single-cell and bulk data. Rather, ScPP connects single-cell with bulk data via the expressions of phenotype-associated marker genes. As a result, ScPP needs not combine single-cell and bulk data into one data for analysis, and thus is more unlikely to confront the serious heterogeneity between single-cell and bulk data. We deem that it is just the simple and clear logic, namely integration of single-cell and bulk data by marker genes instead of by correlations between bulk samples and single cells, to endow ScPP with the merit of fast and effective implementation while excellent and robust performance.

Like Scissor, ScPP involves a parameter (α) that is used to control the number of single cells identified with specific phenotypes. The smaller α is, the less phenotype-associated single cells are inferred, while the accuracy will be higher. In constrast, the larger α will result in more phenotype-associated single cells to be identified, but the accuracy will compromise. Our analysis shows that the performance of ScPP is more stable than that of Scissor with the variation of α. This is significant as users are likely to select different parameters for their tasks.

ScPP is an algorithm that aims to uncover cell subpopulations related to specific phenotypes. Here the phenotype refers to a broad spectrum of features, such as prognosis, drug response, cell type and cell state. Nevertheless, like Scissor and scAB, ScPP is confined in cell subtype identification due to limited cell subtype-associated bulk data available. Thus, to identify cell subtypes, related algorithms could be more advantageous.

## Ethics declarations

## Ethics approval and consent to participate

Not applicable

## Consent for publication

Not applicable

## Availability of data and materials

All the datasets analysed during the current study are publicly available.

## Author’s Contributions

YH performed the research, data analyses and visualization, and wrote the manuscript. RL performed data analyses, visualization, and the R package development. LZ supervised the work. XW conceived this study, designed the algorithm and analysis strategies, and wrote the manuscript. All the authors read and approved the final manuscript.

## Supplementary Information

**Table S1. A summary of the datasets analyzed in this study.**

**Table S2. The phenotype information of bulk samples.**

**Table S3. The marker genes of proliferation and stemness signatures.**

**Fig. S1. Kaplan-Meier curves to compare overall and disease-free survival time between LUAD patients with high versus low expressions of marker genes of LUAD+ and LUAD-cells.** Many of the genes upregulated (red color) in LUAD+ cells are negatively correlated with prognosis of LUAD patients, and many of the genes downregulated (blue color) are positively correlated with prognosis of LUAD patients. The log-rank test *P* values are shown.

**Fig. S2.** LUAD-BS marker genes shows significant positive expression correlations with overall survival in GSE11969 and TCGA-LUAD. The log-rank test *P* values are shown.

