## Supplementary material for "Inferring phenotypes of single cells based on the expression profiles of phenotype-associated marker genes in bulks and single cells": Fig. S1

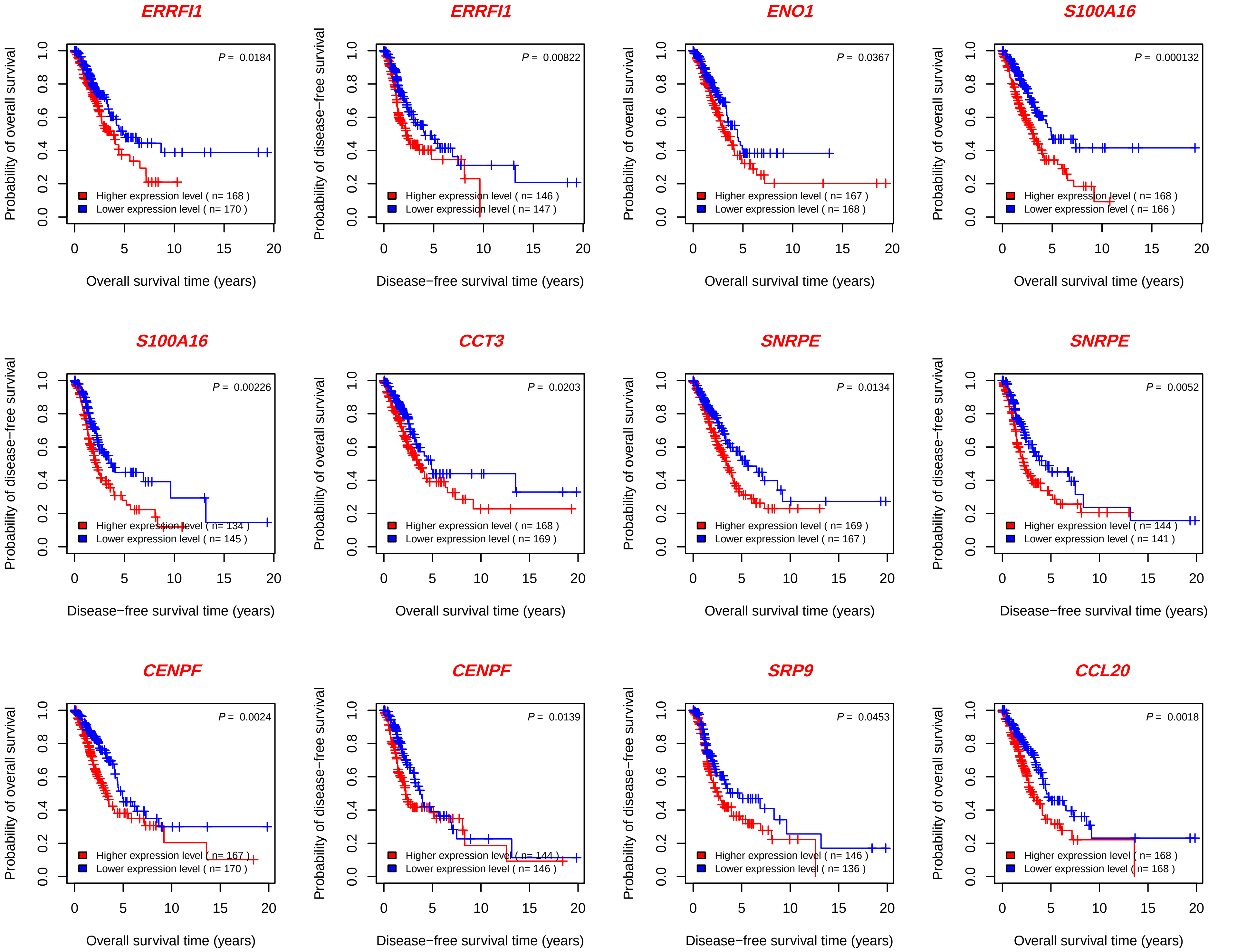
ERRFI1
Probability of overall survival
P = 0.0184
Higher expression level ( n= 168 )
Lower expression level ( n= 170 )
Overall survival time (years)
ERRFI1
Probability of disease−free survival
P = 0.00822
Higher expression level ( n= 146 )
Lower expression level ( n= 147 )
Disease−free survival time (years)
ENO1
Probability of overall survival
P = 0.0367
Higher expression level ( n= 167 )
Lower expression level ( n= 168 )
Overall survival time (years)
S100A16
Probability of overall survival
P = 0.000132
Higher expression level ( n= 168 )
Lower expression level ( n= 166 )
Overall survival time (years)
S100A16
Probability of disease−free survival
P = 0.00226
Higher expression level ( n= 134 )
Lower expression level ( n= 145 )
Disease−free survival time (years)
CCT3
Probability of overall survival
P = 0.0203
Higher expression level ( n= 168 )
Lower expression level ( n= 169 )
Overall survival time (years)
SNRPE
Probability of overall survival
P = 0.0134
Higher expression level ( n= 169 )
Lower expression level ( n= 167 )
Overall survival time (years)
SNRPE
Probability of disease−free survival
P = 0.0052
Higher expression level ( n= 144 )
Lower expression level ( n= 141 )
Disease−free survival time (years)
CENPF
Probability of overall survival
P = 0.0024
Higher expression level ( n= 167 )
Lower expression level ( n= 170 )
Overall survival time (years)
CENPF
Probability of disease−free survival
P = 0.0139
Higher expression level ( n= 144 )
Lower expression level ( n= 146 )
Disease−free survival time (years)
SRP9
Probability of disease−free survival
P = 0.0453
Higher expression level ( n= 146 )
Lower expression level ( n= 136 )
Disease−free survival time (years)
CCL20
Probability of overall survival
P = 0.0018
Higher expression level ( n= 168 )
Lower expression level ( n= 168 )
Overall survival time (years)

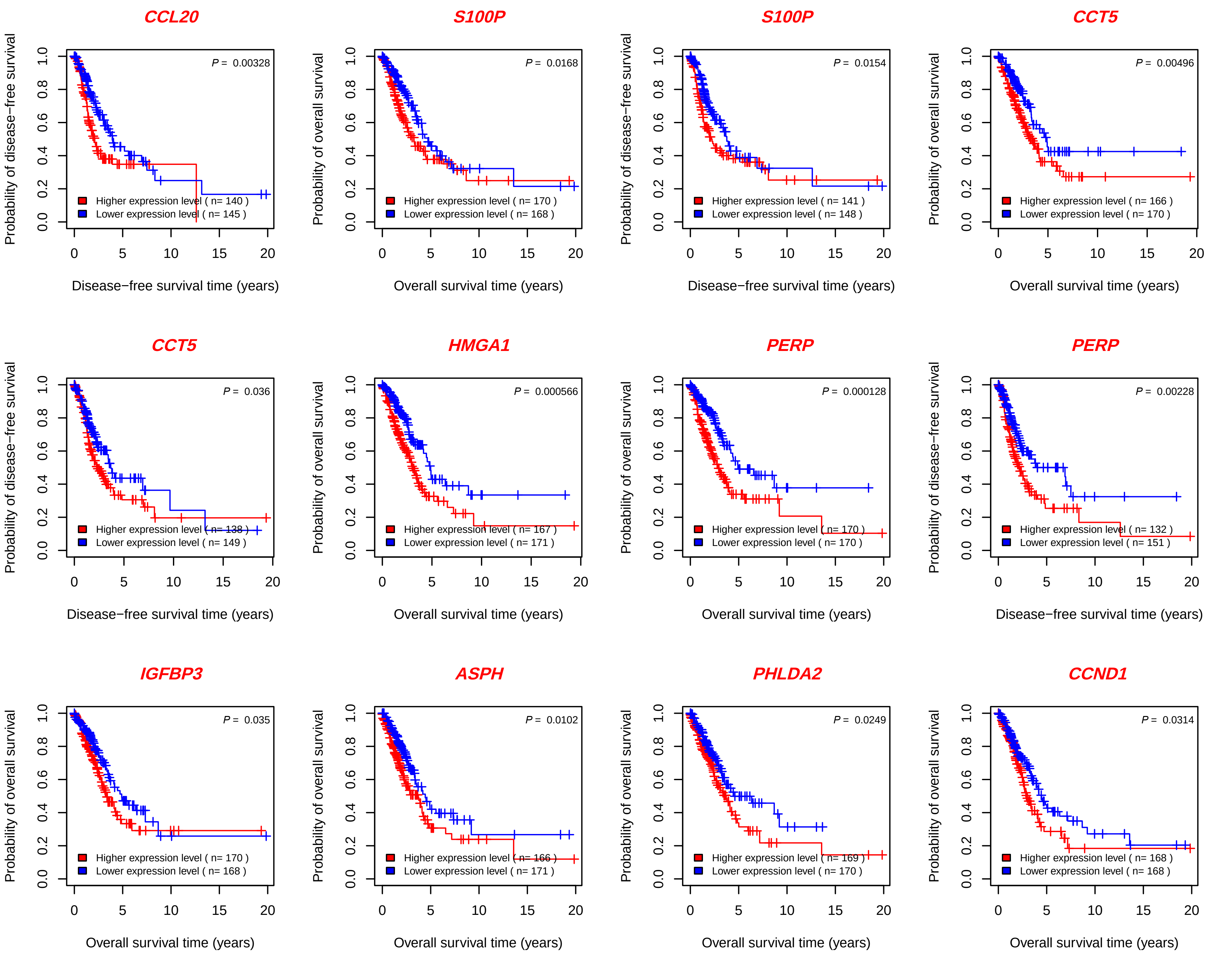
CCL20
P = 0.00328
Higher expression level ( n= 140 )
Lower expression level ( n= 145 )
Probability of disease−free survival
Disease−free survival time (years)
S100P
P = 0.0168
Higher expression level ( n= 170 )
Lower expression level ( n= 168 )
Probability of overall survival
Overall survival time (years)
S100P
P = 0.0154
Higher expression level ( n= 141 )
Lower expression level ( n= 148 )
Probability of disease−free survival
Disease−free survival time (years)
CCT5
P = 0.00496
Higher expression level ( n= 166 )
Lower expression level ( n= 170 )
Probability of overall survival
Overall survival time (years)
CCT5
P = 0.036
Higher expression level ( n= 138 )
Lower expression level ( n= 149 )
Probability of disease−free survival
Disease−free survival time (years)
HMGA1
P = 0.000566
Higher expression level ( n= 167 )
Lower expression level ( n= 171 )
Probability of overall survival
Overall survival time (years)
PERP
P = 0.000128
Higher expression level ( n= 170 )
Lower expression level ( n= 170 )
Probability of overall survival
Overall survival time (years)
PERP
P = 0.00228
Higher expression level ( n= 132 )
Lower expression level ( n= 151 )
Probability of disease−free survival
Disease−free survival time (years)
IGFBP3
P = 0.035
Higher expression level ( n= 170 )
Lower expression level ( n= 168 )
Probability of overall survival
Overall survival time (years)
ASPH
P = 0.0102
Higher expression level ( n= 166 )
Lower expression level ( n= 171 )
Probability of overall survival
Overall survival time (years)
PHLDA2
P = 0.0249
Higher expression level ( n= 169 )
Lower expression level ( n= 170 )
Probability of overall survival
Overall survival time (years)
CCND1
P = 0.0314
Higher expression level ( n= 168 )
Lower expression level ( n= 168 )
Probability of overall survival
Overall survival time (years)

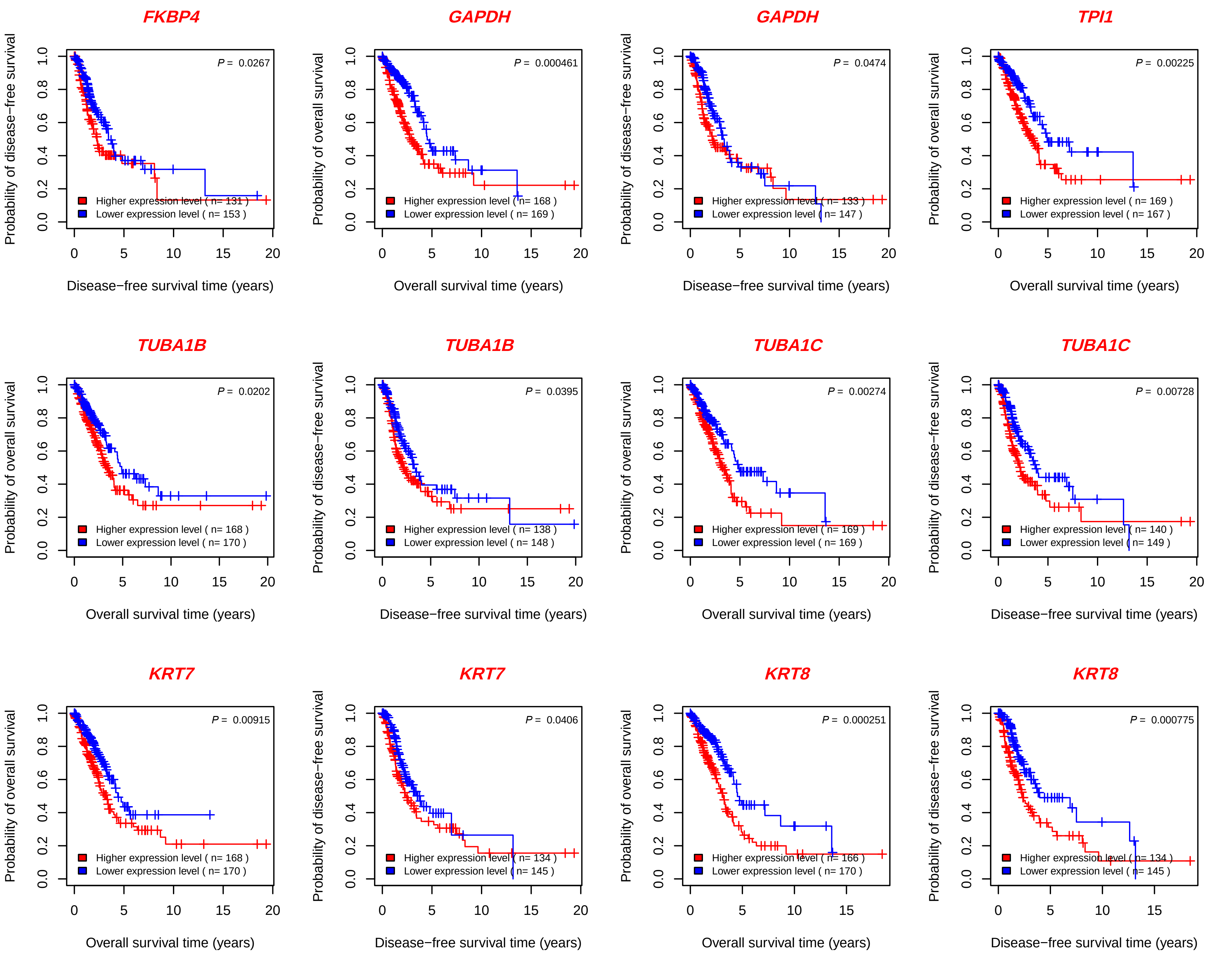

FKBP4
P = 0.0267
Higher expression level ( n= 131 )
Lower expression level ( n= 153 )
Probability of disease−free survival
Disease−free survival time (years)
GAPDH
P = 0.000461
Higher expression level ( n= 168 )
Lower expression level ( n= 169 )
Probability of overall survival
Overall survival time (years)
GAPDH
P = 0.0474
Higher expression level ( n= 133 )
Lower expression level ( n= 147 )
Probability of disease−free survival
Disease−free survival time (years)
TPI1
P = 0.00225
Higher expression level ( n= 169 )
Lower expression level ( n= 167 )
Probability of overall survival
Overall survival time (years)
TUBA1B
P = 0.0202
Higher expression level ( n= 168 )
Lower expression level ( n= 170 )
Probability of overall survival
Overall survival time (years)
TUBA1B
P = 0.0395
Higher expression level ( n= 138 )
Lower expression level ( n= 148 )
Probability of disease−free survival
Disease−free survival time (years)
TUBA1C
P = 0.00274
Higher expression level ( n= 169 )
Lower expression level ( n= 169 )
Probability of overall survival
Overall survival time (years)
TUBA1C
P = 0.00728
Higher expression level ( n= 140 )
Lower expression level ( n= 149 )
Probability of disease−free survival
Disease−free survival time (years)
KRT7
P = 0.00915
Higher expression level ( n= 168 )
Lower expression level ( n= 170 )
Probability of overall survival
Overall survival time (years)
KRT7
P = 0.0406
Higher expression level ( n= 134 )
Lower expression level ( n= 145 )
Probability of disease−free survival
Disease−free survival time (years)
KRT8
P = 0.000251
Higher expression level ( n= 166 )
Lower expression level ( n= 170 )
Probability of overall survival
Overall survival time (years)
KRT8
P = 0.000775
Higher expression level ( n= 134 )
Lower expression level ( n= 145 )
Probability of disease−free survival
Disease−free survival time (years)

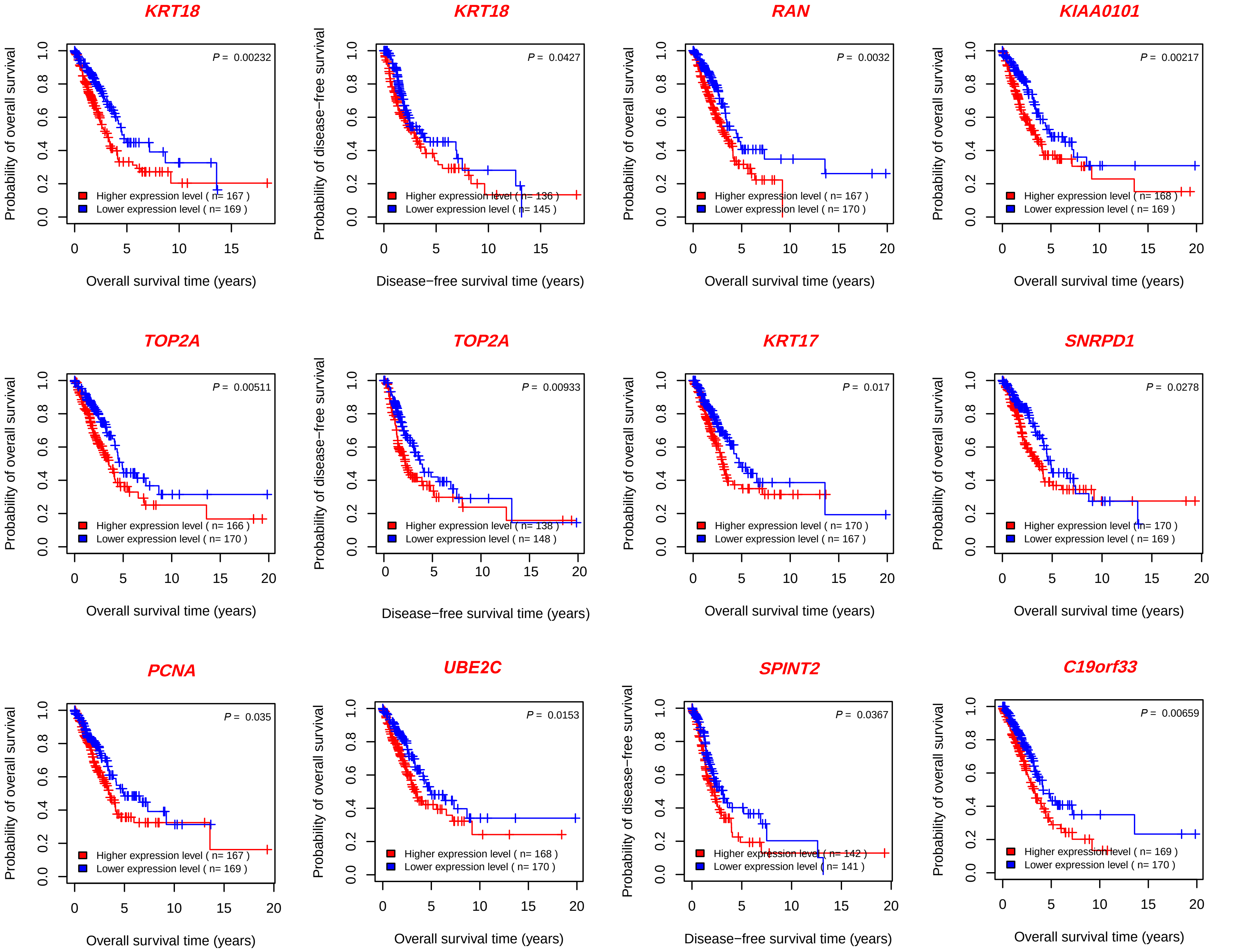
KRT18
P = 0.00232
Higher expression level ( n= 167 )
Lower expression level ( n= 169 )
Probability of overall survival
Overall survival time (years)
KRT18
P = 0.0427
Higher expression level ( n= 136 )
Lower expression level ( n= 145 )
Probability of disease−free survival
Disease−free survival time (years)
RAN
P = 0.0032
Higher expression level ( n= 167 )
Lower expression level ( n= 170 )
Probability of overall survival
Overall survival time (years)
KIAA0101
P = 0.00217
Higher expression level ( n= 168 )
Lower expression level ( n= 169 )
Probability of overall survival
Overall survival time (years)
TOP2A
P = 0.00511
Higher expression level ( n= 166 )
Lower expression level ( n= 170 )
Probability of overall survival
Overall survival time (years)
TOP2A
P = 0.00933
Higher expression level ( n= 138 )
Lower expression level ( n= 148 )
Probability of disease−free survival
Disease−free survival time (years)
KRT17
P = 0.017
Higher expression level ( n= 170 )
Lower expression level ( n= 167 )
Probability of overall survival
Overall survival time (years)
SNRPD1
P = 0.0278
Higher expression level ( n= 170 )
Lower expression level ( n= 169 )
Probability of overall survival
Overall survival time (years)
PCNA
P = 0.035
Higher expression level ( n= 167 )
Lower expression level ( n= 169 )
Probability of overall survival
Overall survival time (years)
UBE2C
P = 0.0153
Higher expression level ( n= 168 )
Lower expression level ( n= 170 )
Probability of overall survival
Overall survival time (years)
SPINT2
P = 0.0367
Higher expression level ( n= 142 )
Lower expression level ( n= 141 )
Probability of disease−free survival
Disease−free survival time (years)
C19orf33
P = 0.00659
Higher expression level ( n= 169 )
Lower expression level ( n= 170 )
Probability of overall survival
Overall survival time (years)

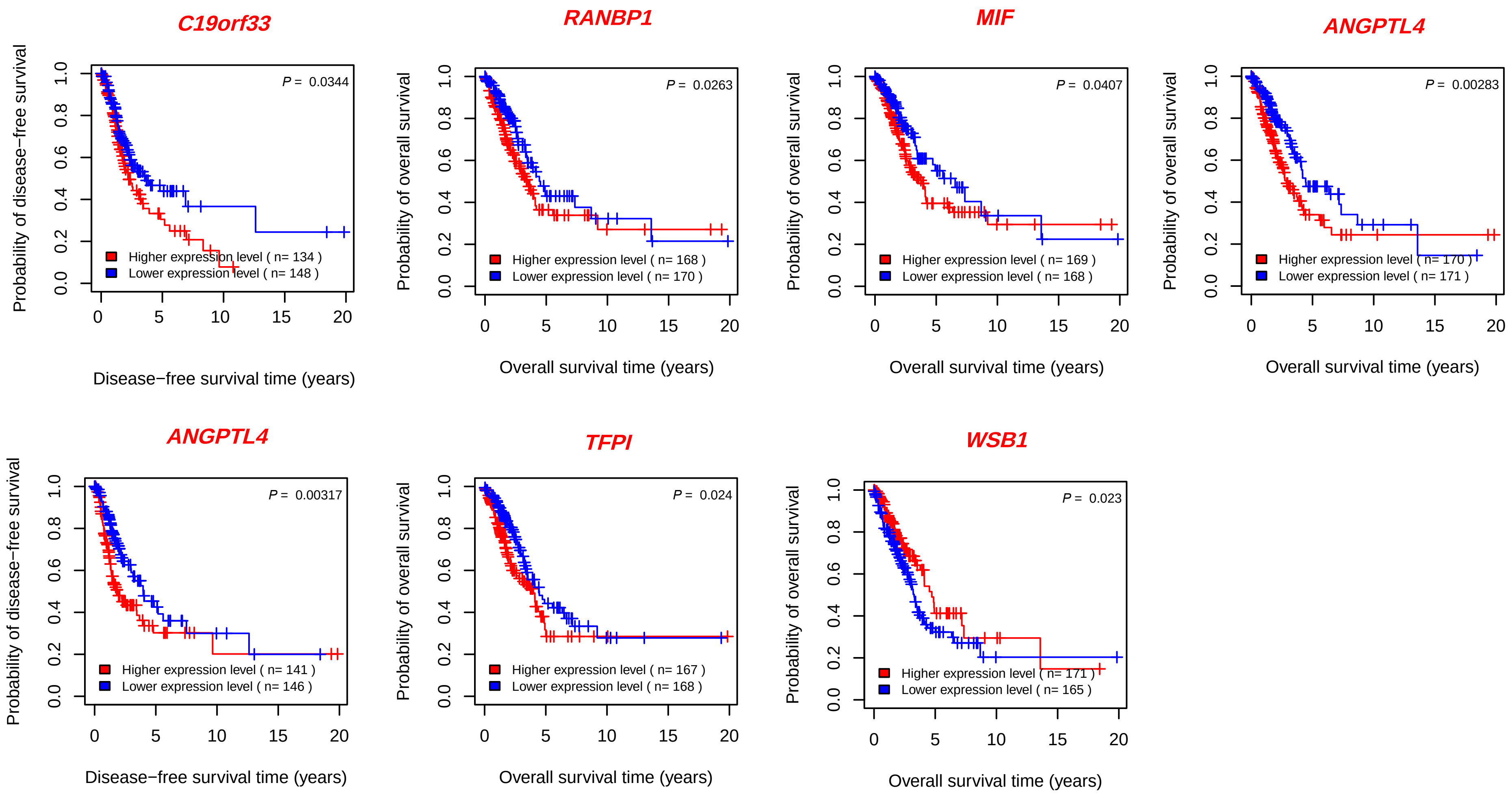
C19orf33
P = 0.0344
Higher expression level ( n= 134 )
Lower expression level ( n= 148 )
Probability of disease−free survival
Disease−free survival time (years)
RANBP1
P = 0.0263
Higher expression level ( n= 168 )
Lower expression level ( n= 170 )
Probability of overall survival
Overall survival time (years)
MIF
P = 0.0407
Higher expression level ( n= 169 )
Lower expression level ( n= 168 )
Probability of overall survival
Overall survival time (years)
ANGPTL4
P = 0.00283
Higher expression level ( n= 170 )
Lower expression level ( n= 171 )
Probability of overall survival
Overall survival time (years)
ANGPTL4
P = 0.00317
Higher expression level ( n= 141 )
Lower expression level ( n= 146 )
Probability of disease−free survival
Disease−free survival time (years)
TFPI
P = 0.024
Higher expression level ( n= 167 )
Lower expression level ( n= 168 )
Probability of overall survival
Overall survival time (years)
WSB1
P = 0.023
Higher expression level ( n= 171 )
Lower expression level ( n= 165 )
Probability of overall survival
Overall survival time (years)

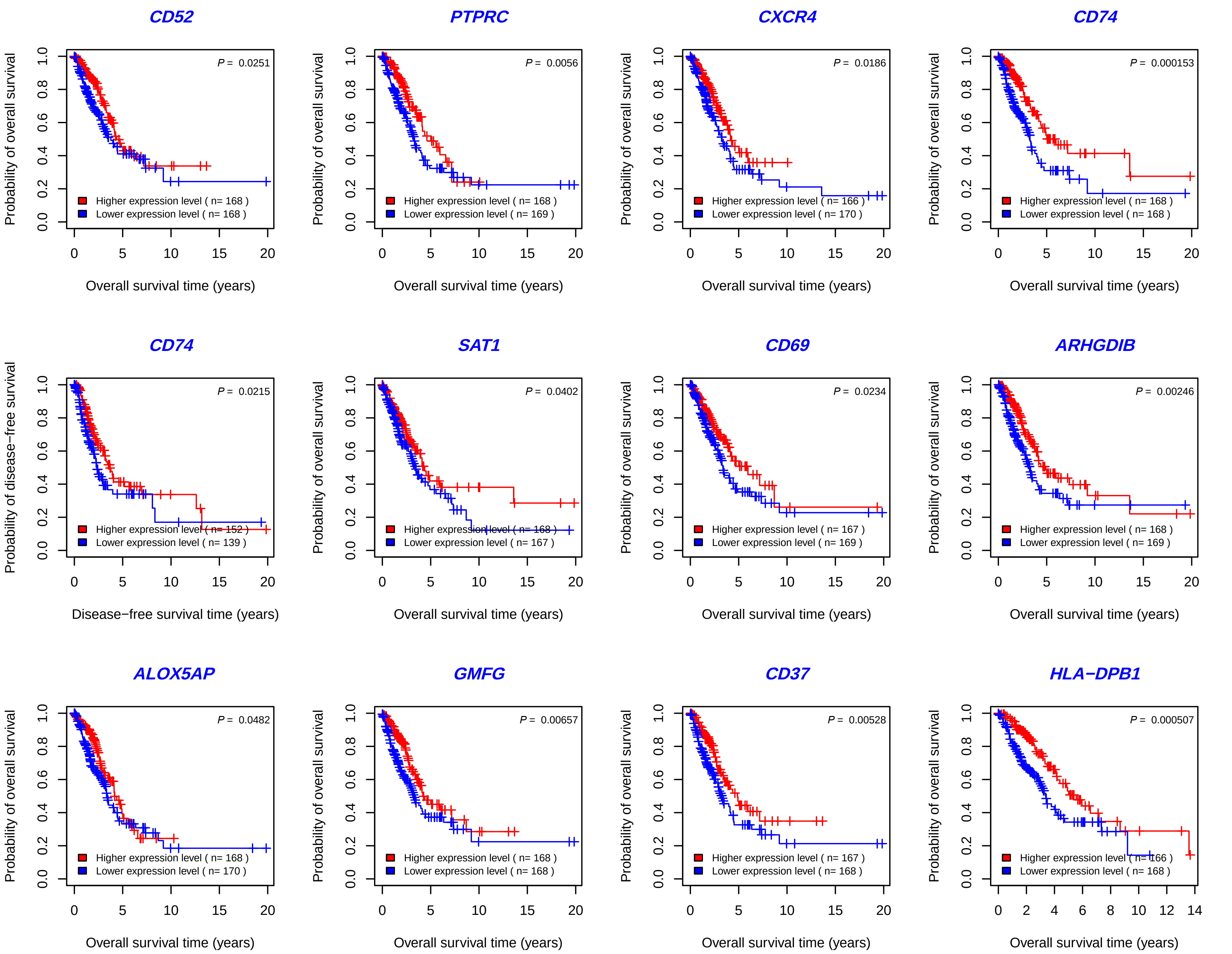
CD52
P = 0.0251
Higher expression level ( n= 168 )
Lower expression level ( n= 168 )
Probability of overall survival
Overall survival time (years)
PTPRC
P = 0.0056
Higher expression level ( n= 168 )
Lower expression level ( n= 169 )
Probability of overall survival
Overall survival time (years)
CXCR4
P = 0.0186
Higher expression level ( n= 166 )
Lower expression level ( n= 170 )
Probability of overall survival
Overall survival time (years)
CD74
P = 0.000153
Higher expression level ( n= 168 )
Lower expression level ( n= 168 )
Probability of overall survival
Overall survival time (years)
CD74
P = 0.0215
Higher expression level ( n= 152 )
Lower expression level ( n= 139 )
Probability of disease−free survival
Disease−free survival time (years)
SAT1
P = 0.0402
Higher expression level ( n= 168 )
Lower expression level ( n= 167 )
Probability of overall survival
Overall survival time (years)
CD69
P = 0.0234
Higher expression level ( n= 167 )
Lower expression level ( n= 169 )
Probability of overall survival
Overall survival time (years)
ARHGDIB
P = 0.00246
Higher expression level ( n= 168 )
Lower expression level ( n= 169 )
Probability of overall survival
Overall survival time (years)
ALOX5AP
P = 0.0482
Higher expression level ( n= 168 )
Lower expression level ( n= 170 )
Probability of overall survival
Overall survival time (years)
GMFG
P = 0.00657
Higher expression level ( n= 168 )
Lower expression level ( n= 168 )
Probability of overall survival
Overall survival time (years)
CD37
P = 0.00528
Higher expression level ( n= 167 )
Lower expression level ( n= 168 )
Probability of overall survival
Overall survival time (years)
HLA−DPB1
P = 0.000507
Higher expression level ( n= 166 )
Lower expression level ( n= 168 )
Probability of overall survival
Overall survival time (years)

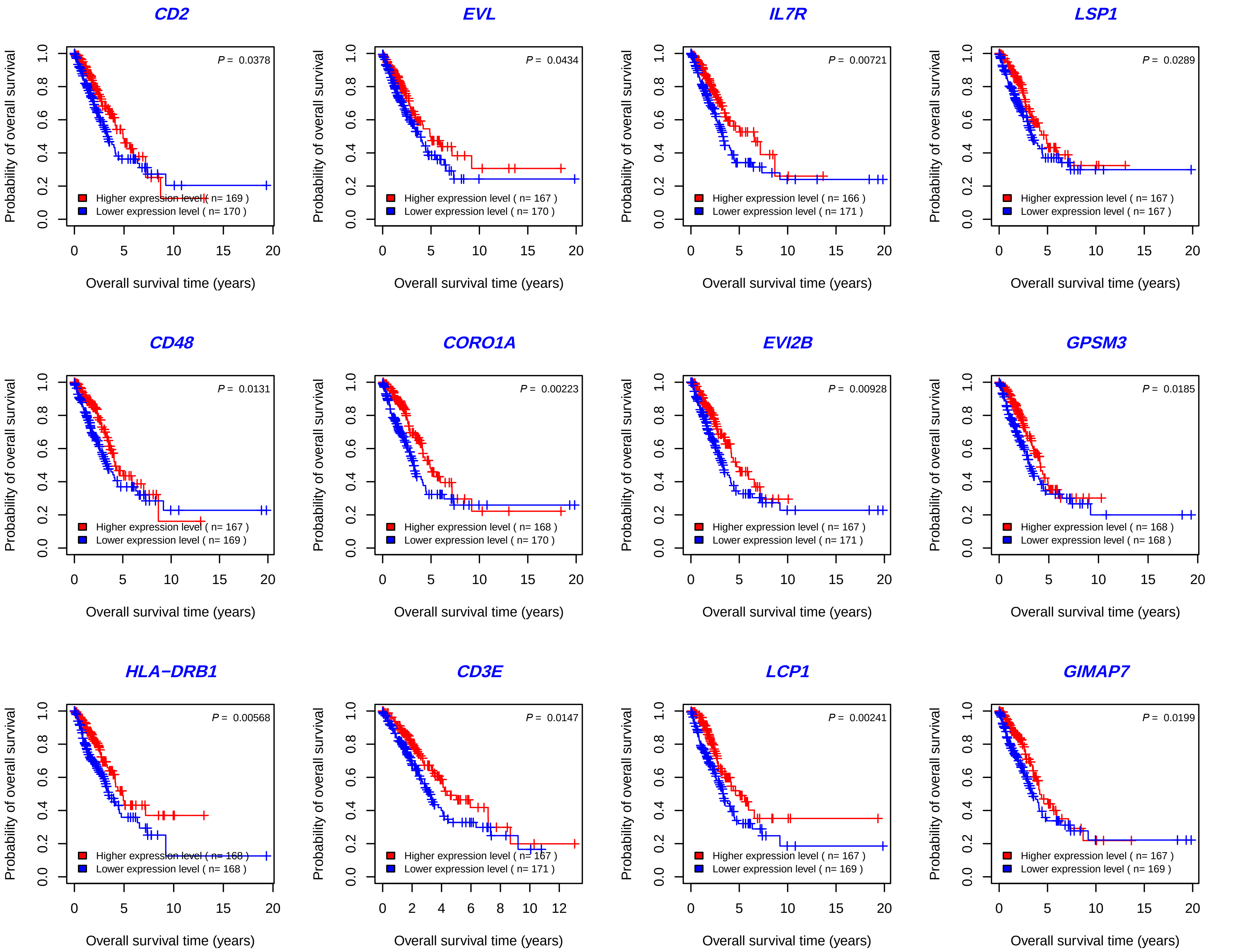
CD2
P = 0.0378
Higher expression level ( n= 169 )
Lower expression level ( n= 170 )
EVL
P = 0.0434
Higher expression level ( n= 167 )
Lower expression level ( n= 170 )
IL7R
P = 0.00721
Higher expression level ( n= 166 )
Lower expression level ( n= 171 )
LSP1
P = 0.0289
Higher expression level ( n= 167 )
Lower expression level ( n= 167 )
CD48
P = 0.0131
Higher expression level ( n= 167 )
Lower expression level ( n= 169 )
CORO1A
P = 0.00223
Higher expression level ( n= 168 )
Lower expression level ( n= 170 )
EVI2B
P = 0.00928
Higher expression level ( n= 167 )
Lower expression level ( n= 171 )
GPSM3
P = 0.0185
Higher expression level ( n= 168 )
Lower expression level ( n= 168 )
HLA−DRB1
P = 0.00568
Higher expression level ( n= 168 )
Lower expression level ( n= 168 )
CD3E
P = 0.0147
Higher expression level ( n= 167 )
Lower expression level ( n= 171 )
LCP1
P = 0.00241
Higher expression level ( n= 167 )
Lower expression level ( n= 169 )
GIMAP7
P = 0.0199
Higher expression level ( n= 167 )
Lower expression level ( n= 169 )
Probability of overall survival
Overall survival time (years)

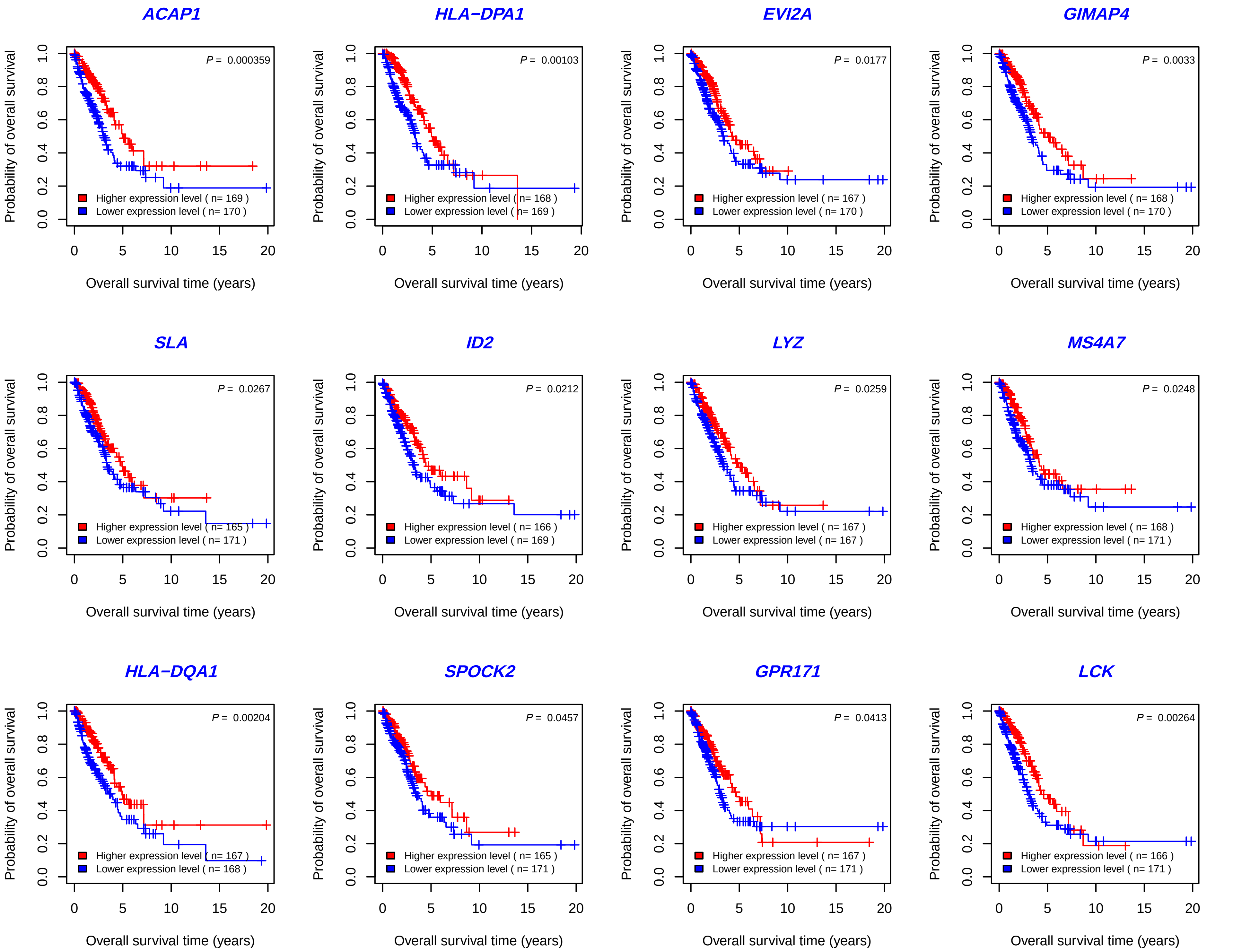
ACAP1
P = 0.000359
Higher expression level ( n= 169 )
Lower expression level ( n= 170 )
HLA−DPA1
P = 0.00103
Higher expression level ( n= 168 )
Lower expression level ( n= 169 )
EVI2A
P = 0.0177
Higher expression level ( n= 167 )
Lower expression level ( n= 170 )
GIMAP4
P = 0.0033
Higher expression level ( n= 168 )
Lower expression level ( n= 170 )
SLA
P = 0.0267
Higher expression level ( n= 165 )
Lower expression level ( n= 171 )
ID2
P = 0.0212
Higher expression level ( n= 166 )
Lower expression level ( n= 169 )
LYZ
P = 0.0259
Higher expression level ( n= 167 )
Lower expression level ( n= 167 )
MS4A7
P = 0.0248
Higher expression level ( n= 168 )
Lower expression level ( n= 171 )
HLA−DQA1
P = 0.00204
Higher expression level ( n= 167 )
Lower expression level ( n= 168 )
SPOCK2
P = 0.0457
Higher expression level ( n= 165 )
Lower expression level ( n= 171 )
GPR171
P = 0.0413
Higher expression level ( n= 167 )
Lower expression level ( n= 171 )
LCK
P = 0.00264
Higher expression level ( n= 166 )
Lower expression level ( n= 171 )
Probability of overall survival
Overall survival time (years)

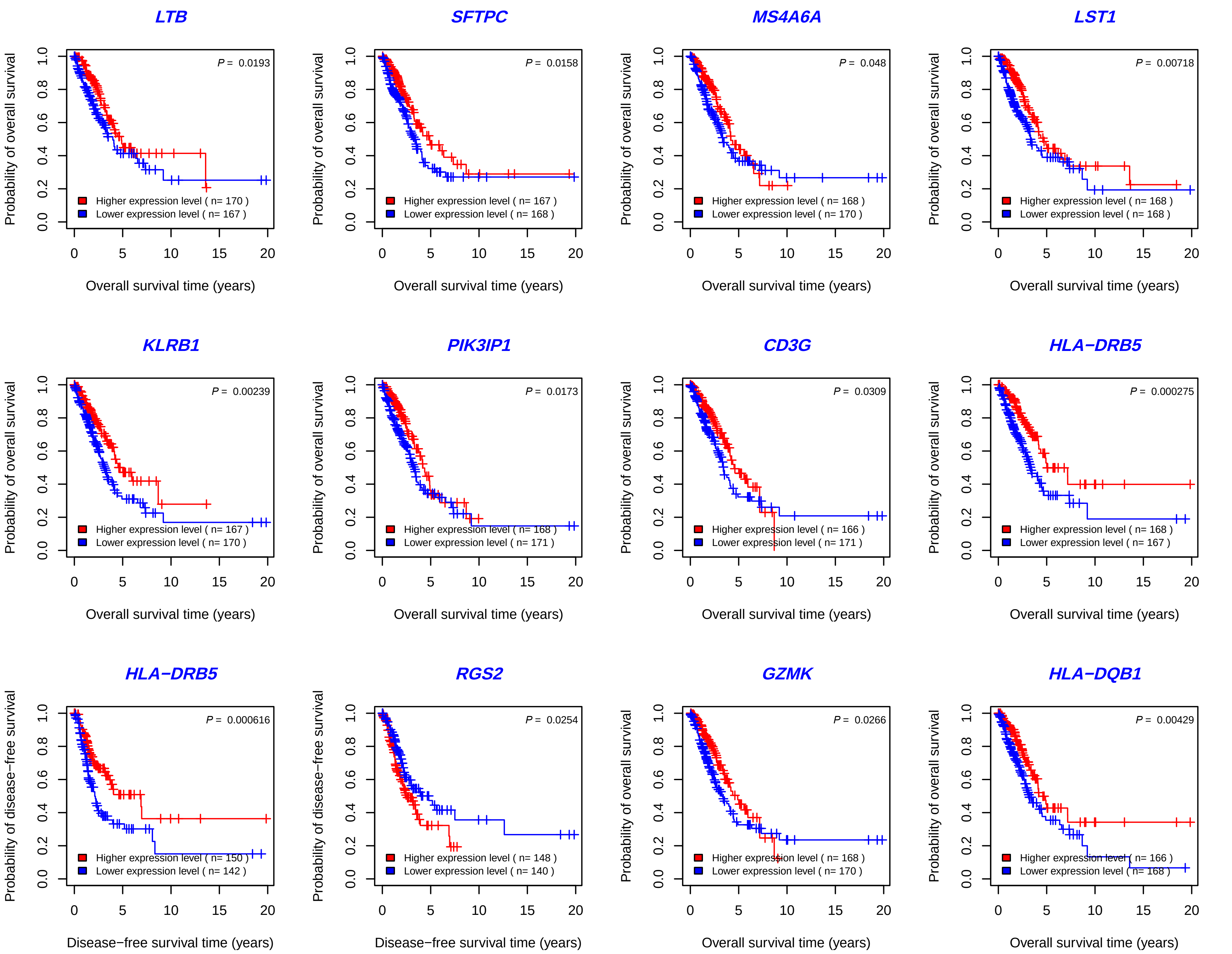
LTB
P = 0.0193
Higher expression level ( n= 170 )
Lower expression level ( n= 167 )
SFTPC
P = 0.0158
Higher expression level ( n= 167 )
Lower expression level ( n= 168 )
MS4A6A
P = 0.048
Higher expression level ( n= 168 )
Lower expression level ( n= 170 )
LST1
P = 0.00718
Higher expression level ( n= 168 )
Lower expression level ( n= 168 )
KLRB1
P = 0.00239
Higher expression level ( n= 167 )
Lower expression level ( n= 170 )
PIK3IP1
P = 0.0173
Higher expression level ( n= 168 )
Lower expression level ( n= 171 )
CD3G
P = 0.0309
Higher expression level ( n= 166 )
Lower expression level ( n= 171 )
HLA−DRB5
P = 0.000275
Higher expression level ( n= 168 )
Lower expression level ( n= 167 )
HLA−DRB5
P = 0.000616
Higher expression level ( n= 150 )
Lower expression level ( n= 142 )
RGS2
P = 0.0254
Higher expression level ( n= 148 )
Lower expression level ( n= 140 )
GZMK
P = 0.0266
Higher expression level ( n= 168 )
Lower expression level ( n= 170 )
HLA−DQB1
P = 0.00429
Higher expression level ( n= 166 )
Lower expression level ( n= 168 )
Probability of overall survival
Overall survival time (years)
Probability of disease−free survival
Disease−free survival time (years)
0.0
0.2
0.4
0.6
0.8
1.0
0
5
10
15
20

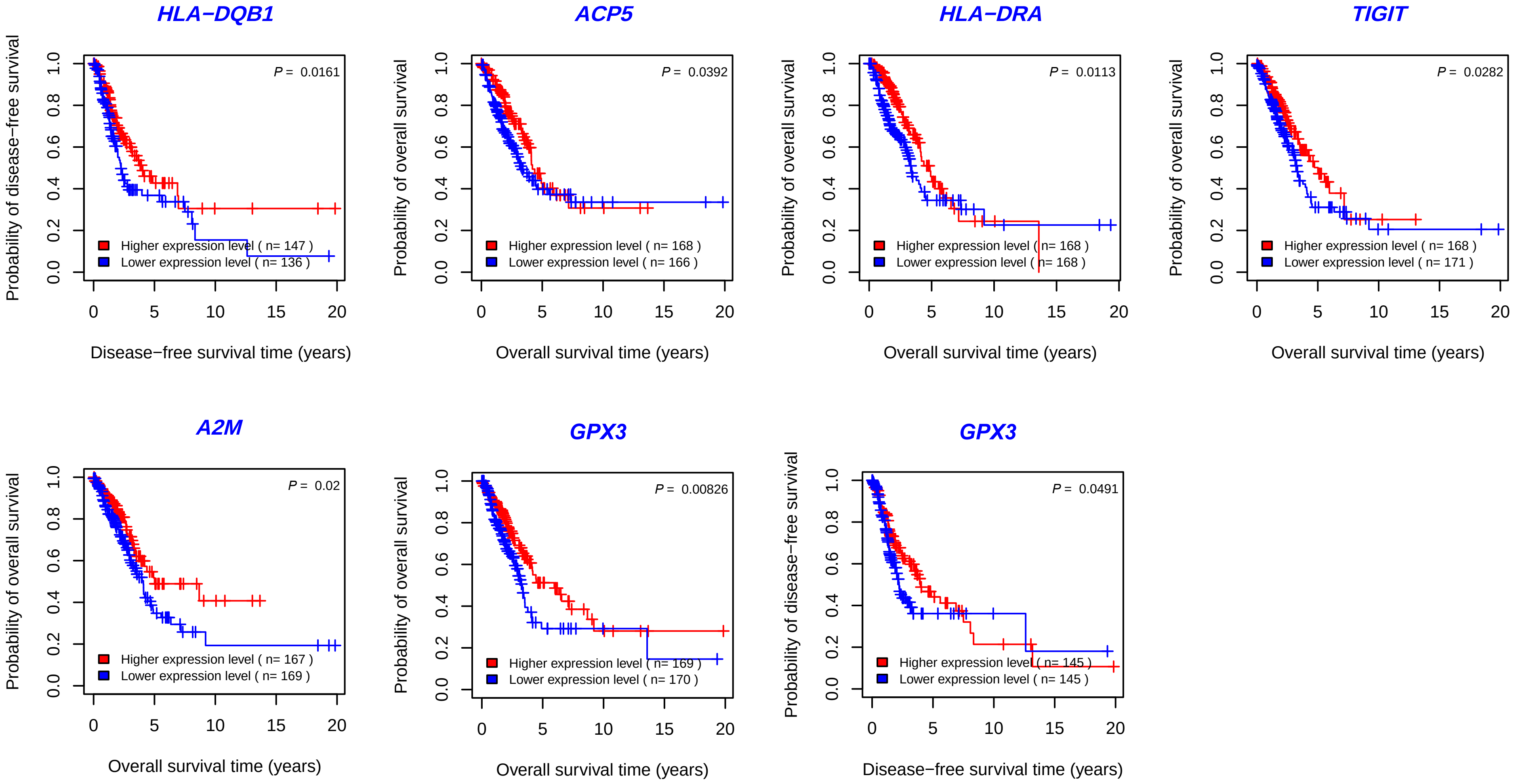


**Fig. S1. Kaplan-Meier curves to compare overall and disease-free survival time between LUAD patients with high versus low expressions of marker genes of LUAD+ and LUAD- cells.** Many of the genes upregulated (red color) in LUAD+ cells are negatively correlated with prognosis of LUAD patients, and many of the genes downregulated (blue color) are positively correlated with prognosis of LUAD patients. The log-rank test P values are shown.
