## Supplementary material for "Inferring phenotypes of single cells based on the expression profiles of phenotype-associated marker genes in bulks and single cells": Fig. S2

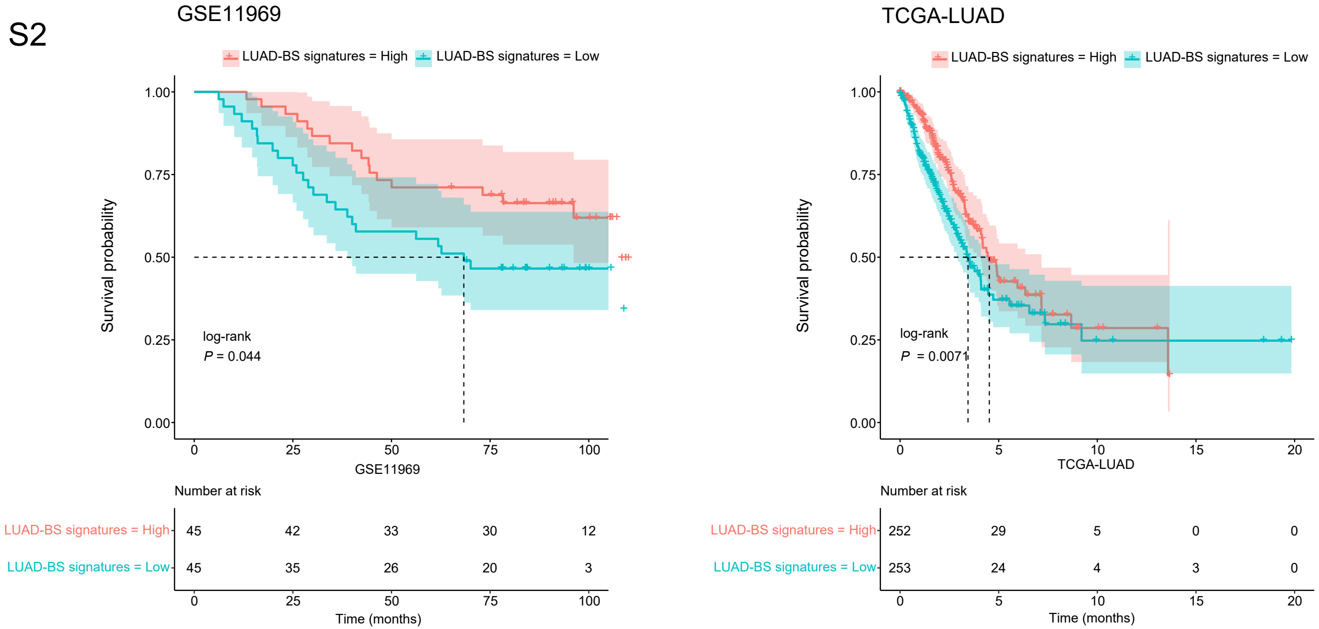


**Fig. S2.** LUAD-BS marker genes shows significant positive expression correlations with overall survival in GSE11969 and TCGA-LUAD. The log-rank test *P* values are shown.
